# Plasma non-esterified fatty acids contribute to increased coagulability in type-2 diabetes through altered plasma zinc speciation

**DOI:** 10.1101/744482

**Authors:** Amélie I. S. Sobczak, Kondwani G. H. Katundu, Fladia A. Phoenix, Siavash Khazaipoul, Ruitao Yu, Fanuel Lampiao, Fiona Stefanowicz, Claudia A. Blindauer, Samantha J. Pitt, Terry K. Smith, Ramzi A. Ajjan, Alan J. Stewart

**Author notes:** Corresponding author: Alan. J. Stewart. School of Medicine, University of St Andrews, Medical and Biological Sciences Building, St Andrews, Fife, KY16 9TF, United Kingdom.

## Abstract

Zn^2+^ is an essential regulator of coagulation and its availability in plasma is fine-tuned through buffering by human serum albumin (HSA). Non-esterified fatty acids (NEFAs) transported by HSA reduce its ability to bind/buffer Zn^2+^. This is important as plasma NEFA levels are elevated in type-2 diabetes mellitus (T2DM) and other diseases with an increased risk of developing thrombotic complications. The presence of 5 mol. eq. of myristate, palmitate, stearate, palmitoleate and palmitelaidate reduced Zn^2+^ binding to HSA. Addition of myristate and Zn^2+^ increased thrombin-induced platelet aggregation in platelet-rich plasma and increased fibrin clot density and clot time in a purified protein system. The concentrations of key saturated (myristate, palmitate, stearate) and monounsaturated (oleate, vaccinate) NEFAs positively correlated with clot density in subjects with T2DM (and controls). Collectively, these data strongly support the concept that elevated NEFA levels contribute to an increased thrombotic risk in T2DM through dysregulation of plasma zinc speciation.

## Introduction

Zinc is an essential modulator of coagulation controlling multiple aspects (1). Molecular regulation of coagulation by zinc is complex, as it binds numerous plasma proteins to influence their activities (1). More specifically, zinc has been shown to enhance platelet aggregation (2), accelerate clotting (3), and delay clot lysis (4). The fraction of zinc responsible for these effects is the “free” aquo ion of Zn^2+^. Low nanomolar free Zn^2+^ is cytotoxic (5, 6); therefore, extracellular zinc is well-buffered. In blood, this buffering role is overwhelmingly performed by human serum albumin (HSA). Approximately 75% of the total 15-20 µM Zn^2+^ is bound to HSA (7), constituting >99% of the labile Zn^2+^ pool (8). Most of the remaining labile Zn^2+^ is bound to small molecules, leaving *∼*1-3 nM free Zn^2+^ under resting conditions (9). However, labile Zn^2+^ concentrations are subject to highly dynamic spatial and temporal variations, increasing sharply during coagulation around activated platelets that release Zn^2+^ from α-granule stores (4, 10). Zn^2+^ is also released by damaged epithelial cells, neutrophils, lymphocytes and erythrocytes and from ruptured atherosclerosis plaques (11).

In addition to binding/buffering Zn^2+^, HSA also transports non-esterified fatty acids (NEFAs) at 5 medium-to high-affinity binding sites (FA1-5), one of which (FA2) is located close to the main Zn^2+^ binding site (site A) (12-14). A secondary Zn^2+^ site with weaker affinity is also present but is unlikely to contribute greatly to Zn^2+^ binding under normal conditions (15). Figure 1A shows the structure of HSA with myristate bound. When a NEFA molecule binds to FA2, the conformation changes, causing site A to be disrupted (the Zn^2+^-coordinating nitrogen of His67 moves ∼8 Å relative to His247 and Asp249; Figure 1B), dramatically reducing the Zn^2+^ affinity of HSA (12-14). Thus, when plasma NEFA levels are elevated, the Zn^2+^ buffering ability of HSA is compromised.

**Figure 1.**
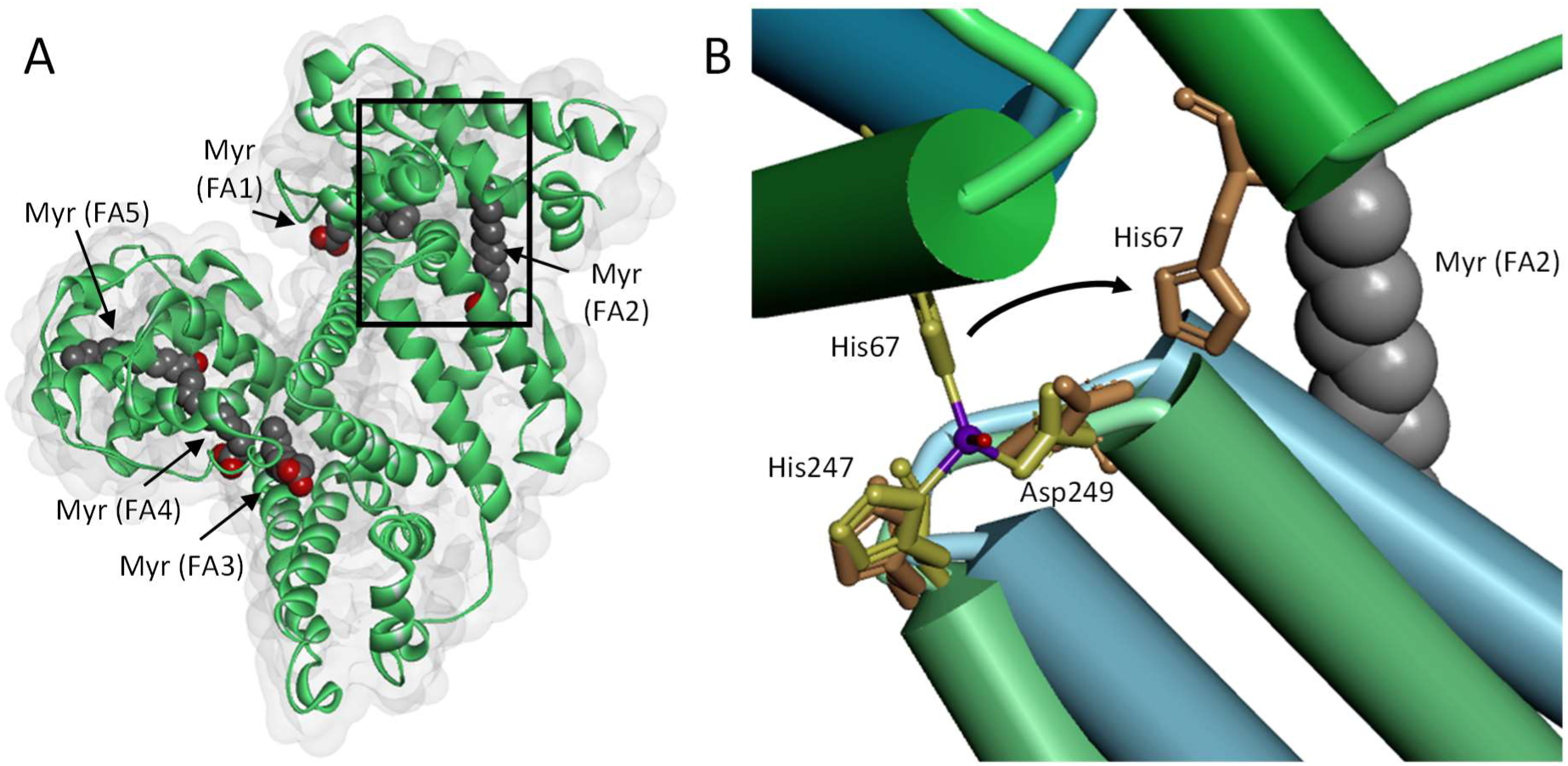
Effects of NEFA binding at the FA2 site on the primary Zn^2+^-binding site of HSA. (A) Structure of HSA with 5 molecules of myristate bound at its high affinity fatty acid binding sites (FA1-5; PDB 1BJ5).(28) The surface of the protein is shown in grey. (B) Overlay of Zn^2+^-binding site in myristate-bound HSA structure (in green, PDB 1BJ5) with the Zn^2+^-bound HSA structure (in blue, PDB 5IJF).(23) The zinc atom is shown in purple and the oxygen from the water participating in Zn^2+^-coordination is in red. A structural change in domain II is induced by the binding of myristate to the FA2 binding site. This triggers the Zn^2+^-coordinating residue His67 to move ∼8 Å away from the site.

Several disease states are associated with elevated plasma NEFA levels including cancer (16), obesity (17), non-alcoholic fatty liver disease (18) and type 2 diabetes mellitus (T2DM) (19). Interestingly, these disorders all associate with an increased risk of thrombotic complications. We hypothesise that high plasma NEFA concentrations disrupt Zn^2+^ binding by HSA, causing more Zn^2+^ to interact with coagulation proteins and consequently enhancing thrombotic risk (20, 21). Here, the ability of plasma NEFA levels to impact on Zn^2+^ handling and blood coagulability is examined. To assess this, Zn^2+^ binding to HSA in the presence of various NEFAs was measured by isothermal titration calorimetry (ITC). The impact of these interactions on platelet aggregation, fibrin clot formation and clot lysis was explored. Moreover, the relationship between plasma NEFA concentration and coagulability in plasma samples taken from individuals with T2DM and controls (without diabetes) was determined. The results support the concept that elevated NEFA levels contribute to an increased thrombotic risk in T2DM through mishandling of plasma Zn^2+^.

## Results

### Influence of various NEFAs on Zn^2+^ binding to HSA

The ability of different NEFAs (octaonate (C8:0), laurate (C12:0), myristate (C14:0), palmitate (C16:0), palmitoleate (C16:1-*cis*), palmitelaidate (C16:1-*trans*) and stearate (C18:0)) to influence Zn^2+^ binding to HSA was assessed by ITC. Building on previous work (13, 15), a “two sets-of-sites” model was chosen to monitor changes to site A stoichiometry in the presence of NEFAs. Fitted isotherms are shown in Figure 2, raw data in Figures S1-25, fitting parameters in Table S1 and fitting results in Table S2. Addition of up to 5 mol. eq. of octanoate had little effect on Zn^2+^ binding to HSA, but a change was seen with laurate and longer chain saturated NEFAs, where the fitting results suggested a reduction in the stoichiometry of site A with increasing NEFA concentration. Indeed, 4 and 5 mol. eq. of laurate reduced site A availability to 0.24 and 0.00, respectively whereas 3 mol. eq. had little effect. In the presence of myristate, palmitate and stearate, 3-5 mol. eq. of these NEFAs reduced binding of Zn^2+^ to HSA in a concentration-dependent manner; 3 mol. eq. myristate or palmitate reduced site A availability to about 50%, and 4 mol. eq. palmitate or stearate sufficient to abolish binding at site A. Almost no Zn^2+^ binding was observed at site A in the presence of 5 mol. eq. of myristate, palmitate or stearate. The unsaturated NEFA, palmitelaidate led to reduced Zn^2+^ binding in a similar manner to palmitate while palmitoleate had less effect than palmitate at the concentrations examined, with the exception of 5 mol. eq. of NEFA where almost no binding was detected.

**Figure 2.**
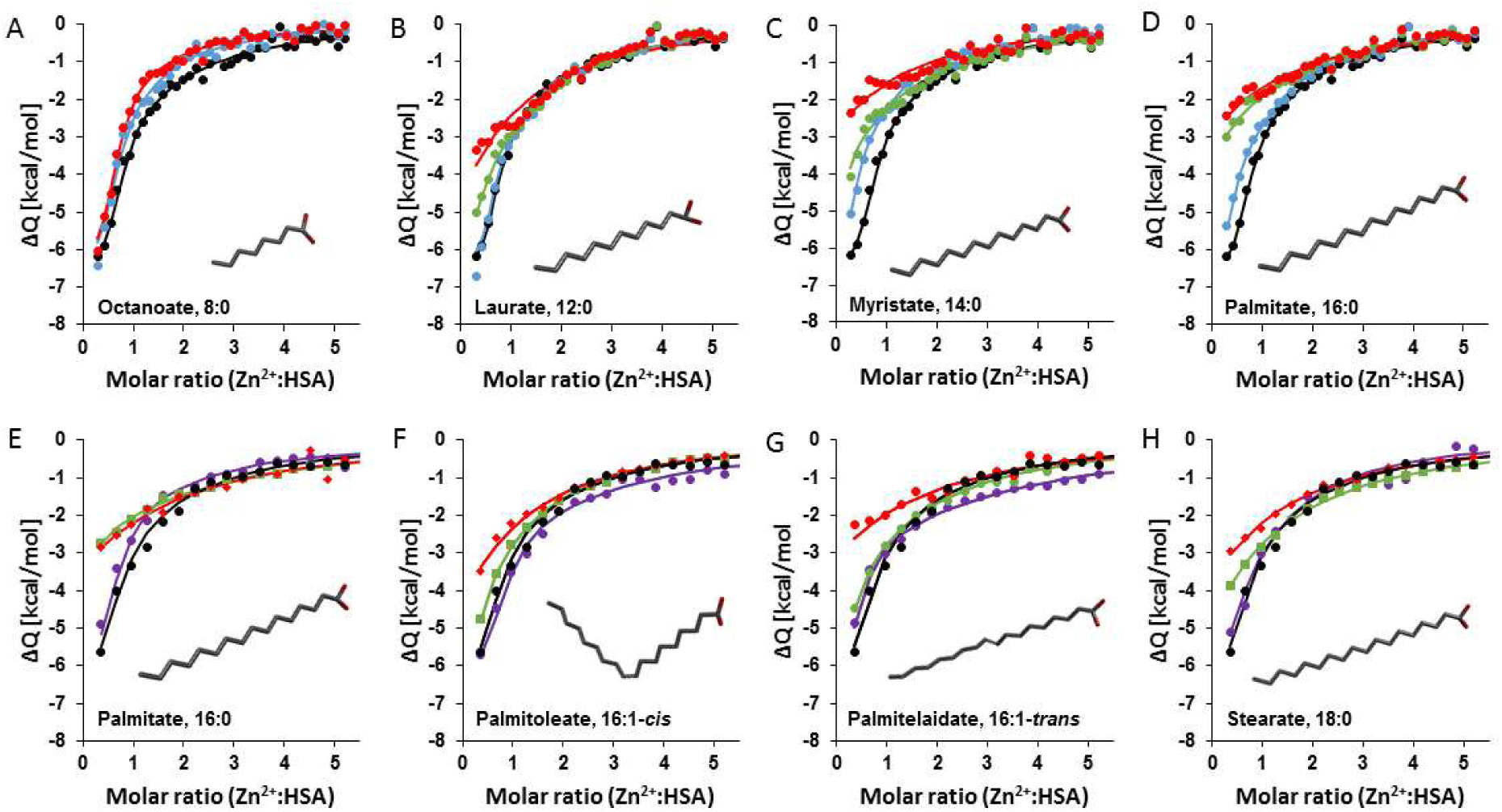
Binding of Zn^2+^ to HSA in presence of different NEFAs. ITC experiments were conducted in a buffer containing 50 mM Tris, 140 mM NaCl, pH 7.4. A first set of experiments were performed with 1.5 mM ZnCl_2_ titrated into 60 µM HSA, in the presence of either 0 (black), 2.5 (purple), 3 (blue), 4 (green) or 5 (red) mol. eq. of different NEFAs: (A) octanoate, (B) laurate, (C) myristate and (D) palmitate. The settings used were 25°C, 39 injections (first injection was 0.4 µL, the remaining injections were 1 µL), initial delay 60 s and spacing 120 s. A second set of experiments were performed with 750 µM Zn^2+^ titrated into 25 µM HSA, in the presence of (E) palmitate, (F) palmitoleate, (G) palmitelaidate and (H) stearate. For those ITC experiments, the settings were adjusted to 25°C, 19 injections (first injection was 0.4 µL, the remaining injections were 2 µL), initial delay 60 s and spacing 120 s. Each fit corresponds to a two-sets-of-sites model. All NEFAs except octanoate perturbed Zn^2+^ binding to the protein, with the effect increasing with the concentration of NEFAs.

### Effect of Zn^2+^ and NEFAs on platelet aggregation

The effect of Zn^2+^ and NEFAs on platelet aggregation in washed-platelets and platelets-in-plasma was assessed (Figure 3 and S26). In washed-platelets, addition Zn^2+^ increased maximum aggregation (p=0.0080, as measured with one-way ANOVA), with a trend observed at 50 µM and a significant increase at 100 µM Zn^2+^ (p=0.0566 and p=0.0047 respectively, measured by Dunnet’s multiple comparison tests). In platelets re-suspended in plasma, no difference was observed upon addition of Zn^2+^, presumably due to Zn^2+^ buffering by HSA. The effect of both octanoate and myristate on platelet aggregation was examined to compare effects of NEFAs with differing abilities to perturbs Zn^2+^ binding to HSA (while aware that Zn^2+^ is already present in plasma). Addition of octanoate had no effect on maximum aggregation. In contrast, addition of 4 mol. eq. of myristate increased maximum aggregation (p=0.0006). The zinc-selective chelator, TPEN, abolished the effects of myristate. Finally, we assessed the effect of 100 µM Zn^2+^ in combination with 4 mol. eq. myristate. This further increased maximum aggregation (compared to 4 mol. eq. myristate alone, p=0.0208).

**Figure 3.**
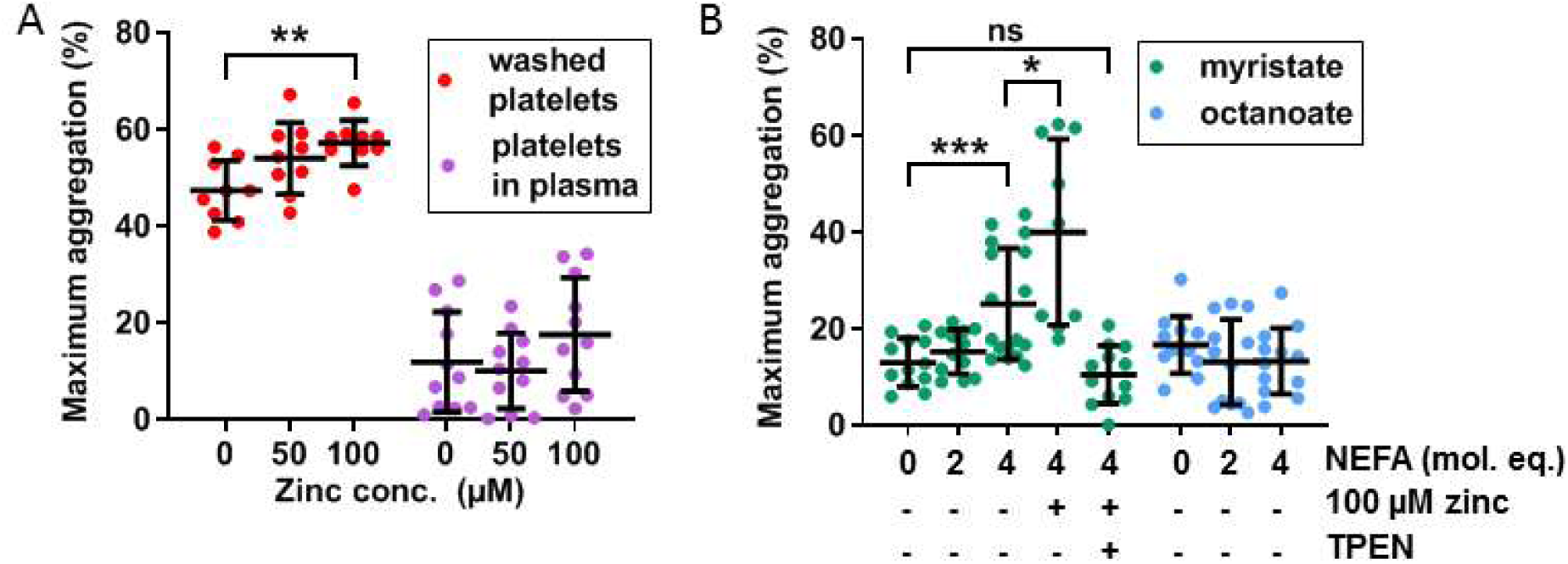
Effects of Zn^2+^ and NEFAs on maximum platelet aggregation and aggregation rate in washed-platelets and platelets re-suspended in plasma. (A) Effect of Zn^2+^ on maximum platelet aggregation (n=11). Maximum aggregation was higher in washed-platelets than in platelets re-suspended in plasma. Addition of Zn^2+^ also increased maximum aggregation in washed-platelets (p=0.0080) but not in platelets re-suspended in plasma. (B) Effect of NEFAs on platelet maximum aggregation in platelets-in-plasma (n=12). Maximum aggregation increased with addition of myristate (p=0.0006) but not octanoate. The presence of 4 mol. eq. myristate and 100 µM Zn^2+^ increases maximum aggregation (n=9, p=0.0208). The addition of the Zn^2+^ chelator TPEN reversed the effect of 4 molecular equivalent of myristate. The data is represented as mean ± SD. Statistical significance is indicated by ns where p>0.05, * where p<0.05, ** where p<0.01 and *** where p<0.001.

### Effect of Zn^2+^ and NEFAs on clot formation and lysis

To examine whether NEFAs alter the effect of Zn^2+^ on fibrin clot formation and lysis, we utilised a validated turbidimetric assay employing both purified protein (fibrinogen and HSA) and plasma samples. In the purified system, addition of 20-100 µM Zn^2+^ significantly increased maximum absorbance and lysis time concentration-dependently, while 20 and 40 µM Zn^2+^ increased clot time and 100 µM Zn^2+^ decreased it (Figure 4 A-C). Zn^2+^ significantly affected all three parameters (p<0.0001 for all). Addition of 4 mol. eq. of myristate also increased maximum absorbance and clot time (p=0.0046 and p=0.0060 respectively; calculated by Sidak’s multiple comparison tests) suggesting a direct effect on these parameters but did not affect lysis time. To highlight the effect of added Zn^2+^, these parameters values were calculated relative to “no Zn^2+^ added” for the samples with/without NEFA (Figure 4 D-F). Addition of 20 µM Zn^2+^ led to a higher maximum absorbance in the presence of myristate and the decrease in clot time at higher Zn^2+^ concentrations (40 and 100 µM) was more pronounced with myristate present. Zn^2+^ did not affect lysis time with or without myristate differently. Myristate significantly affected maximum absorbance and clot time (p=0.0455 and p=0.0194 respectively), but not lysis time. The effect of Zn^2+^ on clot ultrastructure was examined by measuring fibrin fibre thickness using SEM; addition of 20 µM Zn^2+^ significantly increased fibrin fibre diameter (p<0.0001; Figure 4 G, I, S31 and S32).

**Figure 4.**
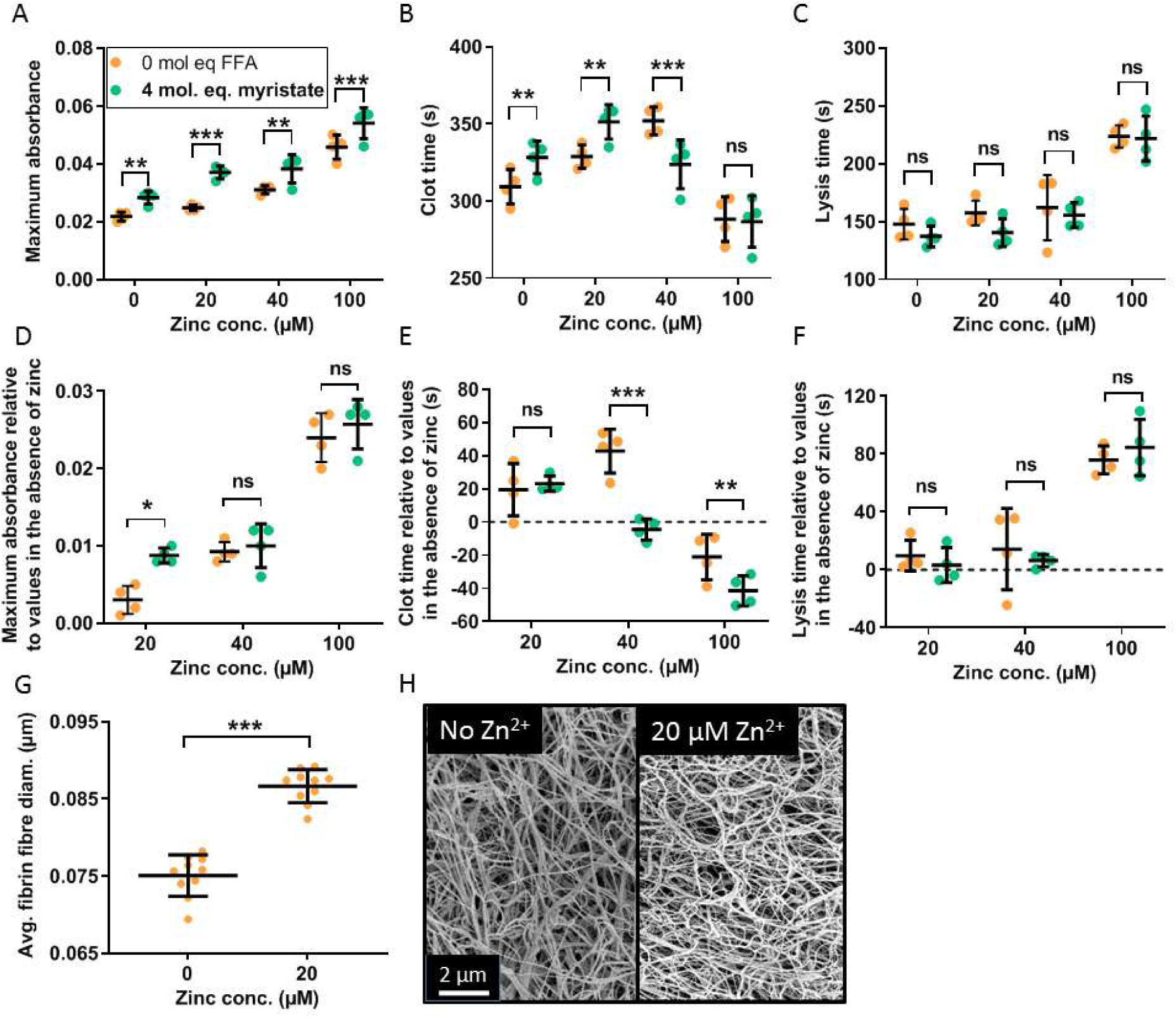
Effects of Zn^2+^ and NEFAs on fibrin clot parameters and on fibrin fibre diameter in a purified system and effects relative to the parameter values in the absence of Zn^2+^. Turbidimetric fibrin clotting and lysis assays were performed in buffer (50 mM Tris, 100 mM NaCl, pH 7.4) with a final concentration of 0.5 mg/mL (2.9 µM) fibrinogen, 100 µM HSA, 2.5 mM CaCl_2_, 0.05 U/mL thrombin, 39 ng/mL tPA, 3.12 µg/mL plasminogen, 0-100 µM ZnCl_2_, and either 0 or 4 mol. eq. myristate (n=4). Fibrin clot parameters including (A) maximum absorbance, (B) clot time and (C) lysis time were measured. Two-way ANOVA followed by Sidak’s multiple comparisons test were used to analyse the data. All three parameters were significantly increased in the presence of Zn^2+^ (p<0.0001, p<0.0001 and p<0.0001 respectively). Maximum absorbance increased in the presence of 4 mol. eq. myristate (p=0.0276), while clot time and lysis time were not significantly altered. The parameter values relative to their values in the absence of Zn^2+^ were then calculated: (D) maximum absorbance, (E) clot time and (F) lysis time. Two-way ANOVA followed by Sidak’s multiple comparisons test were used to analyse the data. All three parameters were significantly increased in the presence of Zn^2+^ (p<0.0001, p<0.0001 and p<0.0001 respectively). Maximum absorbance and clot time were increased in the presence of 4 mol. eq. myristate (p=0.0455 and p=0.0194 respectively), while lysis time was not significantly altered. (G) Fibrin fibre diameters from SEM experiments using a purified system with final concentrations of 4.5 µM fibrinogen, 270 µM HSA, 2.5 mM CaCl_2_, 0.5 U/mL thrombin and either 0 or 18 µM ZnCl_2_ (duplicates of clot, 5 images per samples, 50 fibres measured per images). On the abscise axes, “0 µM Zn^2+”^ refers to no added Zn^2+^ in the system. (H) Representative pictures. Addition of Zn^2+^ significantly increased fibrin fibre diameter (p<0.0001). The data is represented as mean ± SD. Statistical significance is indicated by ns where p>0.05, * where p<0.05, ** where p<0.01 and *** where p<0.001.

For the experiments with citrated pooled plasma, 4-(2-pyridylazo)resorcinol was used to calculate the amount of Zn^2+^ required to obtain the equivalent available Zn^2+^ concentration as in the purified system; this is to account for the Zn^2+^-buffering capacity of citrate. The turbidimetric clot assay was performed on pooled-plasma (Figure S28), where addition of Zn^2+^ increased maximum absorbance and clot time, while 20 µM available Zn^2+^ increased lysis time and 40 and 100 µM available Zn^2+^ decreased it. Zn^2+^ significantly affected all three parameters (p<0.0001 for all). Addition of myristate decreased clot time, increased lysis time (p<0.0001 and p=0.0002 respectively; calculated by Sidak’s multiple comparison tests) and did not affect maximum absorbance. The effects of the addition of Zn^2+^ on maximum absorbance, clot time and lysis time were more pronounced in the presence of myristate. Myristate significantly affected maximum absorbance, clot time and lysis time (p=0.0013, p<0.0001 and p<0.0001 respectively).

### Differences in clot formation in plasma from subjects with T2DM and controls

Elevated plasma NEFA levels are associated with T2DM (19, 22), To explore whether NEFA levels in individuals with T2DM may impact on Zn^2+^ handling and coagulability, we analysed plasma from individuals with T2DM and controls. In each sample, NEFA, total zinc and HSA concentrations were measured (Figure 5 A-C). Demographic information and plasma concentrations of lipids and glucose for the two groups are presented (Table S3). The groups were matched for age but not sex (although no sex difference in NEFA concentration was found, Figure S29); the T2DM group had significantly higher BMI, total plasma NEFA and HSA concentrations than controls (p<0.0001, p=0.0011 and p<0.0001 respectively). The T2DM group had higher concentrations of HbA1c, plasma glucose and triglycerides and a higher cholesterol/LDL ratio (p<0.0001, p<0.0001, p=0.0313 and p=0.0198 respectively), but had lower concentrations of cholesterol, HDL and LDL (p<0.0001 for all). Total zinc and fibrinogen concentrations were comparable between groups.

**Figure 5.**
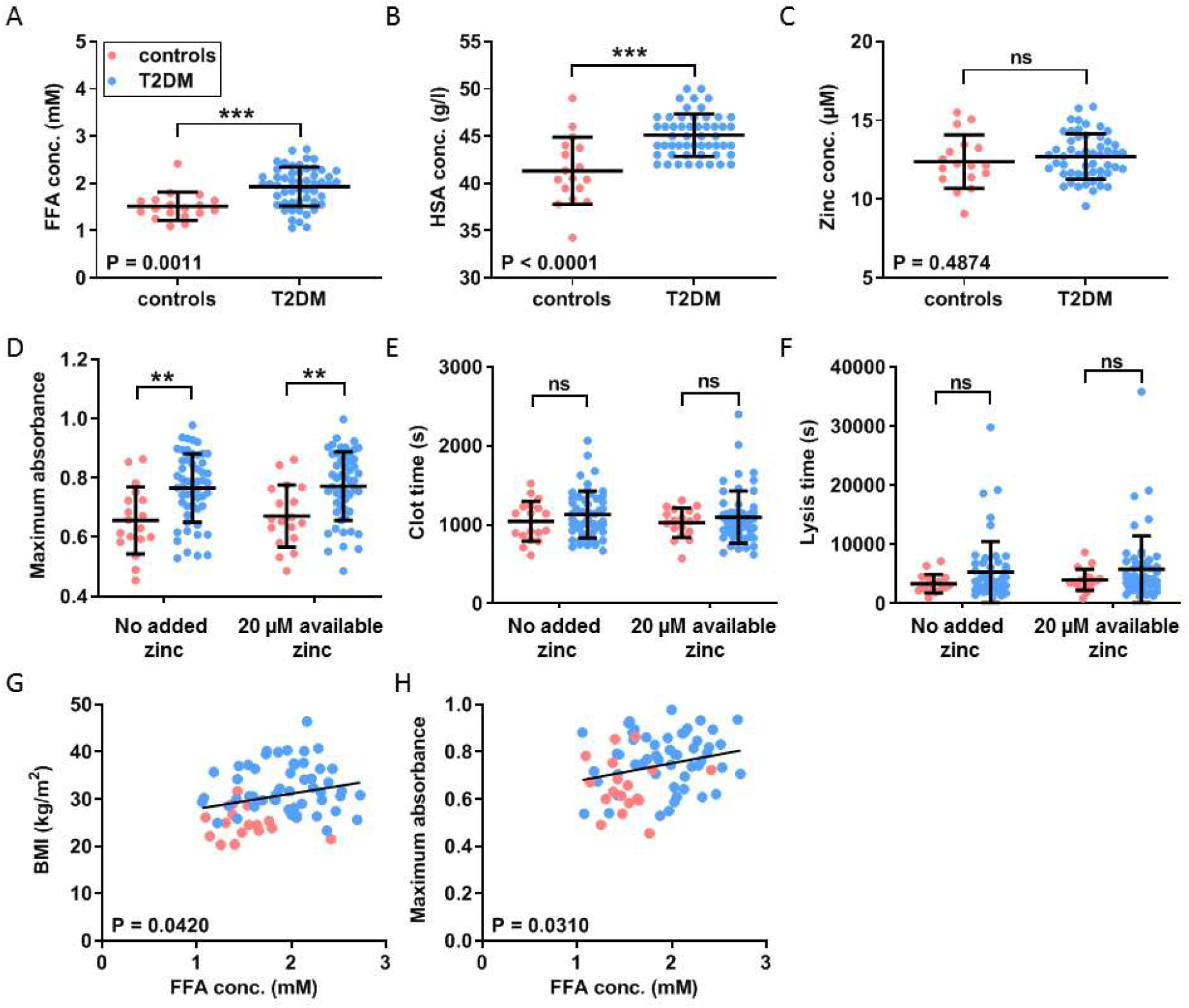
Comparison of the plasma concentrations of NEFA, HSA and zinc and of the clot formation and lysis parameters in plasma samples from patient with T2DM and controls. Comparison of (A) plasma NEFA concentration (B) HSA concentration (C) total plasma zinc concentration in the subjects with T2DM and the controls with t-test. (D) Maximum absorbance and (E) clot time. Turbidimetric fibrin clotting and lysis assays were performed in plasma samples with final concentration of plasma diluted 3-fold in buffer, 7.5 mM CaCl_2_, 0.03 U/mL thrombin and 0 or 20 µM available Zn^2+^ as ZnCl_2_ (concentration calculated before the dilution of plasma) (n=54 for diabetes subjects and 18 for controls). (F) Lysis time; the final concentrations used were plasma diluted 6-fold in buffer, 7.5 mM CaCl_2_, 0.03 U/mL thrombin, 20.8 ng/mL tPA and 0 or 20 µM available Zn^2+^ as ZnCl_2_ (concentration calculated before the dilution of plasma). The data is represented as mean ± SD. Two-way ANOVA followed by Sidak multiple comparison tests were used to analyse the data. Maximum absorbance and lysis time increased significantly in T2DM subjects compared to controls (p<0.0001 and p=0.0448 respectively), but not in the presence of Zn^2+^. Clot time was not significantly altered in the presence or absence of Zn^2+^ or when comparing the T2DM group to the control group. (G) Positive correlation between BMI and NEFA levels (p=0.0420). (H) Positive correlation between maximum absorbance and NEFA levels (p=0.0310). Statistical significance is indicated with ns where p>0.05, * where p<0.05, ** where p<0.01 and *** where p<0.001.

Turbidimetric assays were performed on all samples without added Zn^2+^ and with 20 µM available Zn^2+^, as shown in Figure 5 D-F. Maximum absorbance was higher in the T2DM group compared to controls regardless of the presence or absence of additional Zn^2+^ (p<0.0001; two-way ANOVA). Lysis time was prolonged in T2DM subjects (p=0.0448; two-way ANOVA), but the differences between control samples and those with T2DM at 0 and at 20 µM available Zn^2+^ were not significant (Sidak’s multiple comparison tests). No differences in clot time were observed. Positive correlations between NEFA concentration and both BMI and maximum absorbance were observed (Figure 5 G, H; p=0.0420 and p=0.03010 respectively). There was no correlation between NEFA concentration and either clot time or lysis time. Comparisons of maximum absorbance, clot time and lysis time between sexes revealed no significant differences (Figure S30). SEM studies were performed to examine the fibrin fibre thickness in each group (Figure 6 A-C and S33-36), six plasma samples from each group were randomly selected and pooled. Fibres were found to be significantly thicker in the presence of 20 µM available Zn^2+^ in both groups and were thicker in the T2DM group when compared to controls (p<0.0001 for both).

**Figure 6.**
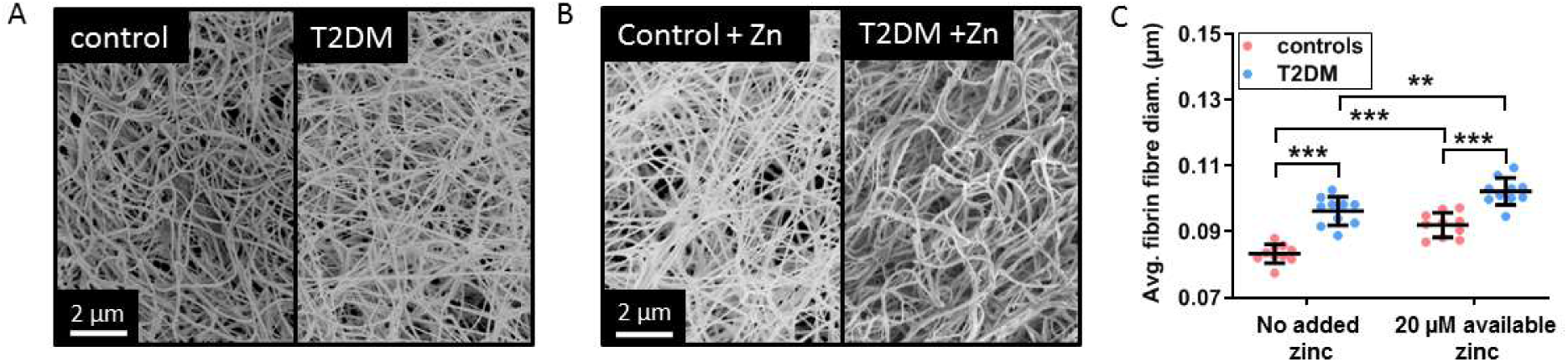
Comparison of the fibrin fibre diameters in plasma samples from patient with diabetes T2DM and controls. SEM experiments were performed using pooled-plasma samples (n=6) from subjects with T2DM and controls with final concentrations of 9:2 plasma in buffer (22.5 µL buffer in a total of 45 µL), 2.5 mM CaCl_2_, 0.5 U/mL thrombin and either 0 or 9 µM available Zn^2+^ as ZnCl_2_ (duplicates of clot, 5 images per sample, 50 fibres measured per image). Representative pictures taken (A) in the absence of Zn^2+^ and (B) in the presence of 9 µM available Zn^2+^. (C) Comparison of fibrin fibre diameter in clots formed from plasma taken from subjects with T2DM and controls. Two-way ANOVA followed by Sidak multiple comparison tests were used to analyse the data. Addition of Zn^2+^ significantly increased fibrin fibre diameter and this effect was exacerbated in the presence of diabetes (p<0.0001 for both). The data is represented as mean ± SD. Statistical significance is indicated with ns where p>0.05, * where p<0.05, ** where p<0.01 and *** where p<0.001.

### Differences in plasma concentrations of specific NEFA species and associations with fibrin clot parameters

The plasma concentration of major NEFA species in the clinical samples was measured using GC-MS (Figure 7). The majority of NEFAs measured were elevated in subjects with T2DM compared to controls (p=0.0010 for myristate, p=0.0023 for palmitate, p=0.0003 for linolenate (18:3), p=0.0054 for oleate (18:1c9), p=0.0029 for vaccinate (18:1c11), p=0.0002 for stearate, p=0.0092 for eicosapentaenoate (20:5) and p=0.0099 for arachidonate (20:4)), with the exception of palmitoleate (16:1), linoleate (18:2), dihomo-γ-linoleate (20:3) and docosahexanoate (22:6) species. The association between plasma concentrations of those NEFAs and fibrin clot maximum absorbance was then assessed. The concentrations of myristate, palmitate, oleate, vaccinate and stearate positively correlated with maximum absorbance (p=0.0313, p=0.0202, p=0.0307, p= 0.0067, p=0.0184 respectively).

**Figure 7.**
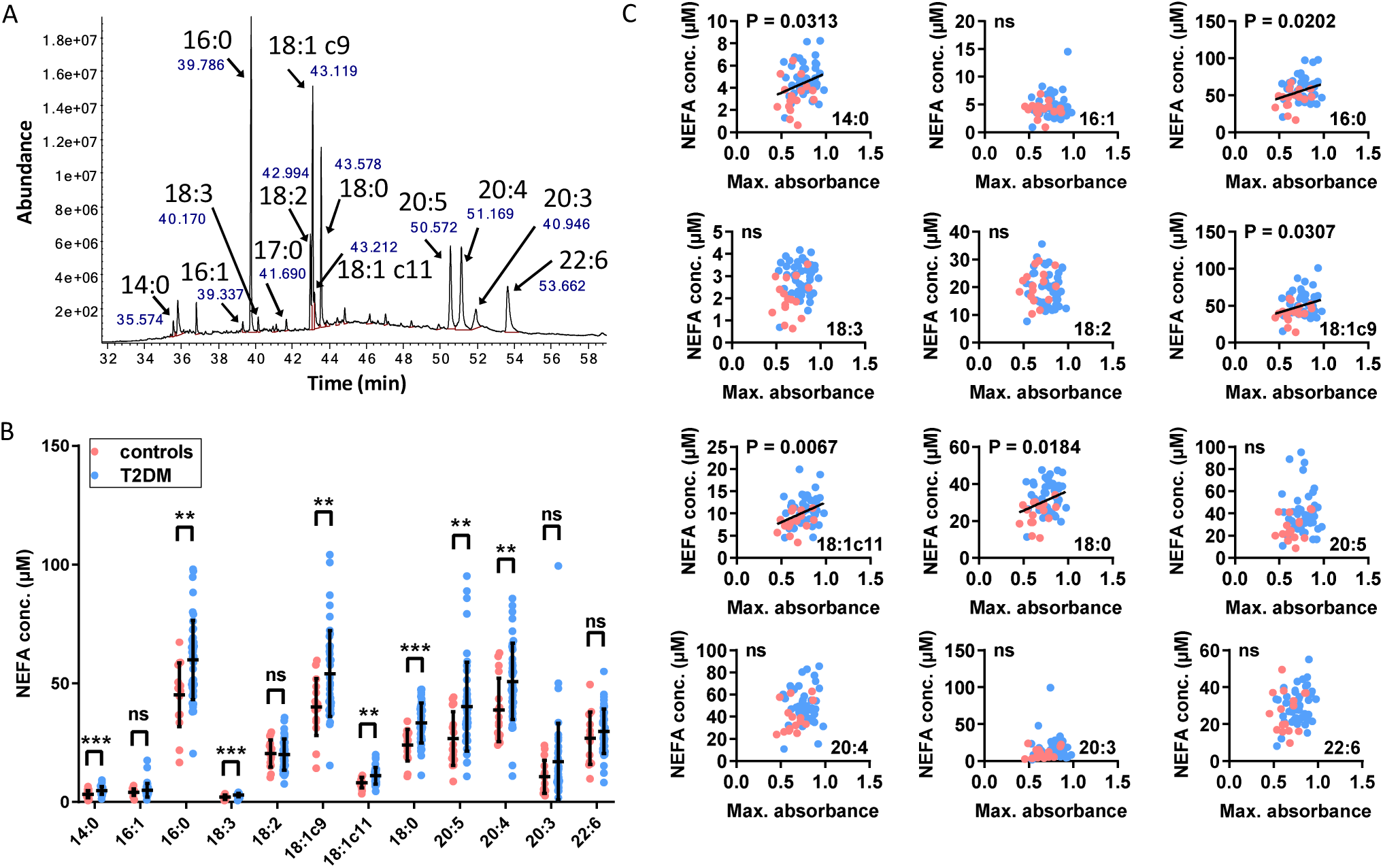
Comparison of the plasma concentration of major NEFA species in samples from patient with diabetes T2DM and controls and associations with fibrin clot parameters. **A**. Example of typical GC-MS chromatogram showing FAME separation. **B**. Comparison of plasma concentrations of NEFAs between subjects with T2DM and controls. The data is represented as mean ± SD. Statistical significance is indicated with ns where p>0.05, * where p<0.05, ** where p<0.01 and *** where p<0.001. Plasma concentrations of myristate (14:0), palmitate (16:0), linolenate (18:3), oleate (18:1c9), vaccinate (18:c11), stearate (18:0), eicosapentaenoate (20:5) and arachidonate (20:4) were elevated in individuals with T2DM (p=0.0010, p=0.0023, p=0.0003, p=0.0054, p=0.0029, p=0.0002, p=0.0092 and p=0.0099 respectively). Concentrations of palmitoleate (16:1), linoleate (18:2), dihomo-γ-linoleate (20:3) and docosahexanoate (22:6) were unchanged. **C**. Relationships between maximum absorbance and the plasma concentrations of the major NEFA species. Plasma concentration of myristate, palmitate, oleate, vaccinate and stearate positively correlated with maximum absorbance (p=0.0313, p=0.0202, p=0.0307, p= 0.0067, p=0.0184 respectively).

## Discussion

We previously demonstrated that myristate (14:0) impacts upon Zn^2+^ binding to HSA at the highest affinity Zn^2+^ site using ITC (15), whilst mutagenesis and X-ray crystallography studies confirm this to be site A (23). Longer-chain saturated NEFAs, which are more abundant in plasma than myristate (e.g. palmitate and stearate) (24) bind to HSA with higher affinity (25), while unsaturated NEFAs bind HSA at the FA2 site with high affinity as long as the degree of unsaturation is low (26). Here, we assessed the ability of various NEFAs to influence binding of Zn^2+^ to HSA. A concentration of 4 mol. eq. of palmitate or stearate almost completely abrogated Zn^2+^ binding to site A, with 3 mol. eq. also reducing Zn^2+^ binding capacity. Palmitoleate had less of an effect on Zn^2+^ binding compared to palmitate and palmitelaidate (no X-ray crystallographic structure of palmitoleate or palmitelaidate binding to HSA is currently available to explain this phenomenon). The total NEFA concentrations in plasma from subjects with T2DM measured in this work were as high as 2.7 mM, which is equivalent to >4 mol. eq. of total NEFA relative to HSA concentration. Although our binding experiments examined the effects of NEFAs on Zn^2+^-HSA interactions in isolation (whilst in plasma there is a mixture of different NEFAs), it would appear that this allosteric interaction nevertheless has the potential to strongly impact on plasma Zn^2+^ handling *in vivo*: when platelets release Zn^2+^ during coagulation, HSA is likely to buffer/control its action in the vicinity of injury sites.

The importance of Zn^2+^ for platelet aggregation is well established (2), with its effects on platelet behaviour exercised through both extracellular and intracellular interactions (27). Here we found that Zn^2+^ concentration-dependently enhanced maximum aggregation in washed platelets. However, in platelets-in-plasma, where HSA and other zinc-binding molecules can buffer Zn^2+^, addition of up to 100 µM Zn^2+^ (0.17 mol. eq. compared to HSA) had no effect on these parameters. Addition of 4 mol. eq. myristate to platelets-in-plasma resulted in an increase in maximum aggregation, which further increased with added Zn^2+^. The observation that the myristate-mediated effects were abolished by zinc-chelating TPEN (Figure 3B), gives strong support to the conclusion that the myristate-mediated effects are due to loss of Zn^2+^-buffering capacity by HSA. The observation that 4 mol. eq. of the octanoate exerted no observable effect on aggregation of platelets-in-plasma is also consistent with the Zn^2+^-NEFA crosstalk hypothesis, as octanoate is too short to elicit the allosteric switch (28), and does not affect Zn^2+^ binding to HSA (Figure 2A).

The effect of Zn^2+^ on fibrin clot characteristics was previously examined by Henderson *et al* (3, 4), who found that Zn^2+^ accelerated clot formation and increased fibrin fibre thickness and clot porosity. This was suggested to promote clot lysis by allowing increased flow of plasma components inside the clot. However, the same studies showed that Zn^2+^ reduced plasminogen activation, resulting in delayed clot lysis. These studies were performed using a purified system similar to ours or in dialysed plasma. The range of Zn^2+^ concentrations was small, up to 6 or 15 µM. Here, we expand on those results by investigating a wider range of Zn^2+^ concentrations (up to 100 µM), as the maximum total Zn^2+^ concentration that can be reached in proximity of activated platelets, although still unknown, will be much higher than basal physiological Zn^2+^ concentrations. In our pooled-plasma experiments, to avoid losing any smaller molecules, we chose not to dialyse the plasma, but instead to estimate the amount of Zn^2+^ to be added in order to obtain the available Zn^2+^ concentrations desired. Our results indicate that in plasma, maximum absorbance and clot time increase at physiological and higher Zn^2+^ concentrations. The lysis time increased in the presence of 20 µM Zn^2+^ as shown previously (4), however it decreased at higher concentrations (40-100 µM Zn^2+^). Also, we confirmed that fibrin fibre thickness increases in the presence of Zn^2+^ (3). The potential effects of NEFA-binding to HSA on coagulation had not been investigated before, but addition of 4 mol. eq. myristate in a purified system that did not include HSA, increased maximum absorbance and clot time (29). Addition of 4 mol. eq. myristate to plasma (which includes physiological zinc) increased lysis time, decreased clot time and did not affect maximum absorbance. To differentiate the standalone-effect of NEFAs from their effect on Zn^2+^-binding by HSA, we calculated parameter values relative to the values in the absence of added Zn^2+^. In plasma, the addition of 4 mol. eq. myristate results in more pronounced changes in maximum absorbance and clot time when Zn^2+^ is added. This shows that, independently to their own effect, NEFAs influence buffering of Zn^2+^ by HSA to consequently alter Zn^2+^ speciation in plasma.

To confirm that pathophysiological concentrations of NEFAs are present in individuals with T2DM and determine the potential impact of elevated NEFA levels on clot formation, we examined plasma samples taken from individuals with T2DM and controls. In accordance with other studies (19, 22), we found that those with T2DM had significantly elevated plasma NEFAs. The T2DM group also had a higher concentration of plasma HSA. No difference in total zinc concentration was observed between the groups, contrary to some previous reports documenting a small decrease in total plasma zinc in individuals with T2DM (30). In our cohort we observed an increase in fibrin fibre diameter in clots formed from pooled-plasma from subjects with T2DM compared to controls. However, the higher fibrinogen concentrations in plasma from T2DM subjects likely contributed to this. A previous examination of clots formed from fibrinogen purified from individuals with T2DM and controls found the T2DM-derived samples exhibited denser and less porous clots (31). This would likely contribute to the higher maximum absorbance (higher density) and longer lysis time (lower porosity) that we observed. We also demonstrate that NEFA concentration associates with clot maximum absorbance, similar to our observation in the purified and the pooled-plasma systems upon addition of myristate. In addition, an examination of the plasma concentrations of major NEFA species confirmed that most were elevated in individuals with T2DM and showed a positive correlation between maximum absorbance and some NEFA species, in particular saturated NEFAs (myristate, palmitate and stearate) and mono-unsaturated NEFAs (oleate and vaccinate), which although we did not look at those *cis*-unsaturated NEFAs specifically, is generally in accord with our ITC results. We did not find any significant difference in clot parameters between males and females with T2DM. However, other studies found maximum absorbance to be higher in females with T2DM (32).

It thus appears from the correlations between (particularly saturated) NEFA levels and clot maximum absorbance that plasma NEFAs induce changes in the speciation of plasma Zn^2+^ to influence Zn^2+^-dependent clotting. The effects of NEFAs and Zn^2+^ on coagulation demonstrated here are summarised in Figure 8. In other studies individuals with T2DM have been reported to be mildly deficient in zinc (30). This could be due to impaired transport of zinc by HSA, resulting in altered zinc distribution and/or partial clearance of the “excess” zinc. Plasma zinc levels have been negatively correlated with the risk of developing cardiovascular diseases in individuals with T2DM (33). Zinc is important for insulin storage and zinc supplementation in T2DM has beneficial effects (improved insulin and glucose levels and decreased risk of developing T2DM) (34-36). However, while supplementation may increase zinc availability, it may also increase zinc binding to other plasma proteins, including coagulation proteins, with further consequences for thrombotic risk in individuals with T2DM. Heparin, an important anticoagulant, has been shown to be increasingly neutralised in the presence of elevated Zn^2+^ levels (11, 37). Thus, the effect of high plasma levels of NEFA and altered Zn^2+^ speciation may need to be carefully considered when choosing an antithrombotic treatment for individuals with T2DM.

**Figure 8.**
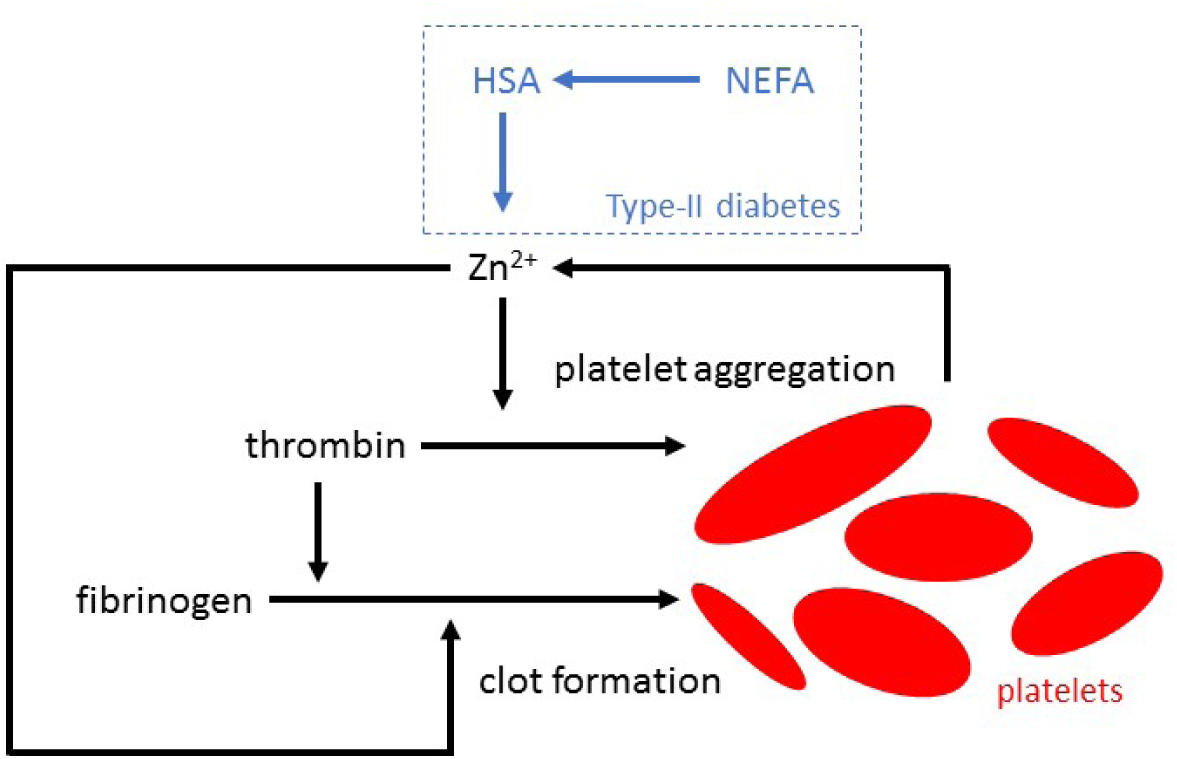
Zn^2+^ increases platelet aggregation and fibrin clot formation. Platelet activation during coagulation triggers the release of Zn^2+^. In addition, NEFA binding to HSA disrupts Zn^2+^ binding at site A. This will likely increase the available plasma Zn^2+^ concentration resulting in an increase in platelets aggregation and influencing fibrin clot formation.

T2DM is not the only condition associated with high plasma NEFAs (22), and although correlations await to be established, it seems prudent to suggest that plasma NEFA levels need to be carefully controlled in all cardiometabolic disorders. This could be achieved through several means, including: **1**. A diet low in saturated fat - the heterogeneity in plasma NEFA levels in our T2DM subjects is likely in part due to diet. In addition, dietary intake of saturated fatty acids and polyunsaturated omega-3 fatty acids have been associated with respectively an increase and a decrease in T2DM occurrence (38-40), however this is now being increasingly challenged (40-45). **2**. The use of statins or fenofibrate as, in addition to affecting cholesterol or triglyceride levels, such drugs have been shown to reduce plasma NEFA levels (46, 47). **3**. Targeting fatty acid synthase in order to reduce *de-novo* synthesis of NEFAs - this could be particularly important in cancer-associated thrombosis, as expression of this enzyme is frequently increased in tumour cells (48). **4**. Design and administration of a small molecule able to inhibit binding of NEFAs to the FA2 site on HSA.

In conclusion, this study shows that plasma NEFA levels correlate with increased coagulability in T2DM. It is also revealed that the NEFA species which exhibit the largest plasma concentration increase in T2DM can perturb zinc binding to HSA and that addition of zinc to plasma induces similar changes in coagulatory functioning as is observed in T2DM. Thus it appears that NEFAs are likely to influence available plasma Zn^2+^ concentrations by a mechanism that involves the FA2 site on HSA, such as to exert a pro-coagulatory effect. Given that plasma NEFA levels can be controlled pharmacologically or via dietary intervention, we believe these findings not only increase our understanding of T2DM but will be useful for the treatment and management of thrombotic complications associated with the disease.

## Materials and Methods

Full details of the methods can be found in the Online Supplementary Appendix.

### Ethical statement

Recruitment of healthy volunteers and blood sample collection for the platelet aggregation study was approved by the School of Medicine Ethics Committee, University of St Andrews. Plasma samples from subjects with T2DM and controls were collected following approval by the National Research Ethics Service Committee Yorkshire & The Humber – Leeds East. All blood samples were taken after obtaining written informed consent.

### Isothermal titration calorimetry

Experiments were performed using a MicroCal iTC200 (Malvern Pananalytical, Malvern, UK) and 50 mM Tris (tris(hydroxymethyl)aminomethane), 140 mM NaCl, pH 7.4 buffer. Two sets of experiments were carried out: (1) titrating 1.5 mM ZnCl_2_ into 60 µM HSA in presence of 0-5 molar equivalents (mol. eq.) of octanoate, laurate, myristate or palmitate; (2) for less soluble FFAs, titrating 0.75 mM ZnCl_2_ into 25 µM HSA in presence of 0-5 mol. eq. of palmitate, palmitoleate, palmitelaidate or stearate. The FFAs were diluted in either methanol or ethanol before being incubated with HSA in the reaction buffer for 2 h at 37°C (1% final alcohol concentration). Heats of dilution were accounted for with blank titrations performed by injecting ligand solution into reaction buffer and subtracting the averaged heat of dilution from the main experiments. Data fitting was performed using AFFINImeter (Santiago de Compostela, Spain). Initial fitting was performed using the Zn^2+^/FFA-free HSA titration and the values obtained were used to fix K1, ΔH1 and N2 for the other titrations.

### Platelet aggregation assays

Platelets were isolated from whole blood collected in acid citrate dextrose collecting tubes from healthy donors recruited from the student body. The blood was spun twice at 23°C, once at 700 × g for 8 min to isolate platelet-rich-plasma and once at 400 × g for 20 min to pellet the platelets. The platelets were washed and re-suspended in buffer solution (145 mM NaCl, 5 mM KCl, 1 mM MgCl_2_, 10 mM HEPES, 1 mM CaCl_2_, 10 mM D-Glucose, pH 7.4) for washed-platelet experiments, or re-suspended in platelet-poor-plasma prepared from hirudin-coated tubes for platelet experiments requiring whole plasma. Hirudin-coated collection tubes were used to avoid chelation of Zn^2+^ by other agents (e.g. citrate or EDTA).

Platelet aggregation experiments were performed to assess the effect of Zn^2+^ and FFAs (octanoate or myristate) on platelet aggregation. Solutions of ZnCl_2_ (10, 50 or 100 µM), sodium octanoate or sodium myristate (2 or 4 mol. eqs.) and N,N,N′,N′-tetrakis(2-pyridinylmethyl)-1,2-ethanediamine (TPEN, 50 µM diluted in ethanol) were added to washed-platelets or platelets-in-plasma. Vehicle control experiments were also performed. Platelet aggregation was elicited with 2 µM of γ-thrombin (Merck, Watford, UK, final volume 200 µL). Absorbance was monitored at 430 nm every 55 s for 35 min using an Optima plate-reader (BMG Labtech, Ortenberg, Germany) while incubating the plate at 37°C and shaking it in orbital mode. Data were recorded as a negative change in absorbance from baseline (0%) and expressed as a percentage of the maximum response (100%). Calibration of 100 % aggregation was achieved using platelet-poor-plasma. From the recorded responses, a maximum aggregation response was obtained.

### Clinical sample collection

A total of 54 patients with T2DM and 18 age-matched controls were recruited from Leeds Teaching Hospital Trust. Individuals with known diagnosis of T2DM and aged between 18-75 years were recruited into the study. Given that aspirin may affect clot structure characteristics (49), all individuals recruited into the study were on aspirin treatment for primary or secondary cardiovascular protection. Exclusion criteria included: any type of diabetes other than T2DM, any coagulation disorder, current or previous history of neoplastic disease, history of acute coronary syndrome or stroke within 3 months of enrolment, active history of transient ischaemic attacks, history of deep venous thrombosis or pulmonary embolism, treatment with oral anticoagulant or non-steroidal anti-inflammatory drugs, abnormal liver function tests defined as alanine transferase >3 fold upper limit of normal, or previous or current history of gastrointestinal pathology. Baseline fasting blood samples were collected in trisodium citrate-or in lithium heparin-coated tubes. Plasma was separated within 2 h of collection by centrifugation at 2,400 × g for 20 min at 4°C, snap frozen in liquid nitrogen and stored at -40°C until analysis.

### Turbidimetric fibrin clotting and lysis assays

Clot assays were performed as previously described (50, 51), using purified proteins, pooled-plasma from controls, as well as plasma samples from subjects with T2DM and age-matched controls. To compensate for the complexation of Zn^2+^ by the citrate present in plasma, 4-(2-pyridylazo)resorcinol was used to generate a calibration curve at 490 nm for different Zn^2+^ concentrations in buffer (50 mM Tris, 100 mM NaCl, pH 7.4) and citrated plasma. The amount of Zn^2+^ that should be added in plasma to yield particular concentrations of available Zn^2+^ was calculated (116 µM in citrated plasma was equivalent to 20 µM in buffer, 232 µM to 40 µM and 580 µM to 100 µM).

Sodium myristate (4 mol. eq.) was incubated for 15 min at 37 °C with HSA (in buffer) or plasma. ZnCl_2_ was added to a final concentration of up to 100 µM Zn^2+^ in the purified system or to available Zn^2+^ concentrations in plasma up to 100 µM. Final concentrations were: (1) In the purified system, 0.5 mg/mL fibrinogen (plasminogen-depleted, Merck), 100 µM HSA, 2.5 mM CaCl_2_, 0.05 U/mL thrombin, 39 ng/mL tissue plasminogen activator (tPA, Technoclone, Vienna, Austria) and 3.12 µg/mL plasminogen (Stratech, Ely, UK). (2) In pooled-plasma (First Link (UK) Ltd, Wolverhampton, UK), plasma diluted 6-fold in buffer, 7.5 mM CaCl_2_, 0.03 U/mL thrombin and 20.8 ng/mL tPA. (3) In plasma from subjects with T2DM for the clot formation assays, plasma diluted 3-fold in buffer, 7.5 mM CaCl_2_ and 0.03 U/mL thrombin. For the clot formation and clot lysis assays, plasma diluted 6-fold in buffer, 3.75 mM CaCl_2_, 0.03 U/mL thrombin and 20.8 ng/mL tPA. The absorbance at 340 nm was read every 12 s at 37°C using a Multiskan FC plate-reader (Thermo Scientific, Paisley, UK). Maximum absorbance, clot time and lysis time were calculated from the raw data.

### Scanning electron microscopy (SEM)

Clots were formed in duplicate from either: 1) 10 µM purified fibrinogen and 300 µM HSA in the presence and absence of 40 µM Zn^2+^. 2) Pooled-plasma from 6 randomly chosen patients with T2DM and age-matched controls in the presence and absence of 20 µM available Zn^2+^. Clots were prepared and processed by stepwise dehydration as previously described (50). All clots were viewed and photographed at ×10,000 magnification using a SU8230 scanning electron microscope (Hitachi, Maidenhead, UK) in 5 different areas. The diameter of 50 fibres per image was measured using Adobe Photoshop (Adobe Systems, San Jose, CA). The mean diameters of each image were used to compare each sample types.

### Measurement of total FFA, HSA and total zinc concentrations

FFAs were extracted from citrated plasma from subjects with T2DM and controls with Dole’s protocol (52). FFA concentrations were measured using the FFA Assay Kit - Quantification (Abcam, Cambridge, UK). Zn^2+^ concentrations were measured in lithium-heparin plasma from subjects with T2DM and controls by inductively coupled plasma-mass spectrometry as described previously (53). HSA levels were measured in heparinised plasma with the bromocresol purple method using an automated analyser (Architect; Abbot Diagnosis, Maidenhead, UK). Plasma concentrations of fibrinogen, high density lipoprotein (HDL), low density lipoprotein (LDL), cholesterol, triglyceride, HbA1c, fasting glucose and platelets were measured with routine methods.

### Measurement of the plasma concentration of specific FFA species by GC-MS

The FFA were characterised and quantified by their conversation to fatty acid methyl esters (FAME) followed by gas chromatography-mass spectrometry analysis. Citrated plasma from subjects with T2DM and controls was thawed and spiked with an internal standard fatty acid C17:0 (100 pM) to allow for normalisation. The FFA were extracted with Dole’s protocol (52). The fatty acids were converted to FAME using 1,500 µl of methanol, 200 µl of toluene and 300 µl of 8% HCl, followed by incubation for 5 hr at 45°C. After cooling, samples were evaporated to dryness with nitrogen. The FAME products were extracted by partitioning between 500 µl of water and 500 µl of hexane and the samples were left to evaporate to dryness in a fume hood. The FAME products were dissolved in 30 µl dichloromethane and 1–2 µl analysed by GC-MS (gas chromatography-mass spectrometry) on a Agilent Technologies (GC-6890N, MS detector-5973) with a ZB-5 column (30 M x 25 mm x 25 mm, Phenomenex), with a temperature program of at 70 °C for 10 min followed by a gradient to 220°C at 5°C /min and held at 220°C for a further 15 min. Mass spectra were acquired from 50-500 amu and the identity of FAMEs was carried out by comparison of the retention time and fragmentation pattern with a various FAME standard mixtures (Supelco) as previously described (54).

### Data analysis and representation

Data are shown as mean ± standard deviation (SD). Graphs were generated and statistical analysis was performed using Prism 7.0 (GraphPad Software, La Jolla, CA). Differences between groups were analysed using multiple Student’s t-tests or analysis of variance (ANOVA) with Dunnet’s or Sidak’s multiple comparisons test, while correlations between linear variants were analysed with Pearson’s correlation. Significance threshold: p≤0.05.

## Acknowledgements

This work was supported by the British Heart Foundation (grant numbers PG/15/9/31270, FS/15/42/31556) and travel grants from the Commonwealth Scholarship Commission (grant number MWCN-2017-294) and the International Co-operation project of Qinghai Province (grant number 2017-HZ-811).

