## Supplementary information for "Plasma non-esterified fatty acids contribute to increased coagulability in type-2 diabetes through altered plasma zinc speciation"

**Table S1. Fitting parameters used for the fitting of the ITC experiments.**

**Table S2. Fitting results from ITC experiments.**

**Figure S1. ITC raw data. Full ITC data (including raw data) for $Zn^{2+}$ binding to HSA in the absence of FFA.**

**Figure S2. ITC raw data. Full ITC data (including raw data) for $Zn^{2+}$ binding to HSA in the presence of 3 mol. eq. of octanoate.**

**Figure S3. ITC raw data. Full ITC data (including raw data) for $Zn^{2+}$ binding to HSA in the presence of 5 mol. eq. of octanoate.**

**Figure S4. ITC raw data. Full ITC data (including raw data) for $Zn^{2+}$ binding to HSA in the presence of 3 mol. eq. of laurate.**

**Figure S5. ITC raw data. Full ITC data (including raw data) for $Zn^{2+}$ binding to HSA in the presence of 4 mol. eq. of laurate.**

**Figure S6. ITC raw data. Full ITC data (including raw data) for $Zn^{2+}$ binding to HSA in the presence of 5 mol. eq. of laurate.**

**Figure S7. ITC raw data. Full ITC data (including raw data) for $Zn^{2+}$ binding to HSA in the presence of 3 mol. eq. of myristate.**

**Figure S8. ITC raw data. Full ITC data (including raw data) for $Zn^{2+}$ binding to HSA in the presence of 4 mol. eq. of myristate.**

**Figure S9. ITC raw data. Full ITC data (including raw data) for $Zn^{2+}$ binding to HSA in the presence of 5 mol. eq. of myristate.**

**Figure S10. ITC raw data. Full ITC data (including raw data) for $Zn^{2+}$ binding to HSA in the presence of 3 mol. eq. of palmitate.**

**Figure S11. ITC raw data. Full ITC data (including raw data) for $Zn^{2+}$ binding to HSA in the presence of 4 mol. eq. of palmitate.**

**Figure S12. ITC raw data. Full ITC data (including raw data) for $Zn^{2+}$ binding to HSA in the presence of 5 mol. eq. of palmitate.**

**Figure S13. ITC raw data. Full ITC data (including raw data) for $Zn^{2+}$ binding to HSA in the absence of FFA.**

**Figure S14. ITC raw data. Full ITC data (including raw data) for $Zn^{2+}$ binding to HSA in the presence of 2.5 mol. eq. of palmitate.**

**Figure S15. ITC raw data. Full ITC data (including raw data) for $Zn^{2+}$ binding to HSA in the presence of 4 mol. eq. of palmitate.**

**Figure S16. ITC raw data. Full ITC data (including raw data) for $Zn^{2+}$ binding to HSA in the presence of 5 mol. eq. of palmitate.**

**Figure S17. ITC raw data. Full ITC data (including raw data) for $Zn^{2+}$ binding to HSA in the presence of 2.5 mol. eq. of palmitoleate.**

**Figure S18. ITC raw data. Full ITC data (including raw data) for $Zn^{2+}$ binding to HSA in the presence of 4 mol. eq. of palmitoleate.**

**Figure S19. ITC raw data. Full ITC data (including raw data) for $Zn^{2+}$ binding to HSA in the presence of 5 mol. eq. of palmitoleate.**

**Figure S20. ITC raw data. Full ITC data (including raw data) for $Zn^{2+}$ binding to HSA in the presence of 2.5 mol. eq. of palmitelaidate.**

**Figure S21. ITC raw data. Full ITC data (including raw data) for $Zn^{2+}$ binding to HSA in the presence of 4mol. eq. of palmitelaidate.**

**Figure S22. ITC raw data. Full ITC data (including raw data) for $Zn^{2+}$ binding to HSA in the presence of 5 mol. eq. of palmitelaidate.**

**Figure S23. ITC raw data. Full ITC data (including raw data) for $Zn^{2+}$ binding to HSA in the presence of 2.5 mol. eq. of stearate.**

**Figure S24. ITC raw data. Full ITC data (including raw data) for $Zn^{2+}$ binding to HSA in the presence of 4 mol. eq. of stearate.**

**Figure S25. ITC raw data. Full ITC data (including raw data) for $Zn^{2+}$ binding to HSA in the presence of 5 mol. eq. of stearate.**

**Figure S26. Representative raw data from platelet aggregation assays.**

**Figure S27. Representative raw data from turbidimetric fibrin clotting and lysis assays.**

**Table S3. Demographic information and the results from plasma analysis for the T2DM and control subjects.**

**Figure S28. Effects of $Zn^{2+}$ and FFAs on fibrin clot parameters in pooled plasma and effects relative to the parameter values in the absence of $Zn^{2+}$.**

**Figure S29. Comparison of FFA concentrations between sexes in plasma samples from patient with T2DM and controls.**

**Figure S30. Comparison of clotting parameters between sexes in plasma samples from patient with T2DM and controls.**

**Figure S31. Representative image from SEM experiments. Purified system, no zinc**

**Figure S32. Representative image from SEM experiments. Purified system, 20 μM zinc**

**Figure S33. Representative image from SEM experiments. Controls, no zinc**

**Figure S34. Representative image from SEM experiments. Controls, 20 μM zinc**

**Figure S35. Representative image from SEM experiments. T2DM, no zinc**

**Figure S36. Representative image from SEM experiments. T2DM, 20 μM zinc**

**Table S1. Fitting parameters used for the fitting of the ITC experiments.** ITC data fitting approaches for ITC experiments examining $Zn^{2+}$-binding to HSA in the presence of different FFA. The values for K1' and ΔH1 in the fit "FFA present" are derived from the fit "no FFA" (see Table S2). All entries marked "v" signify parameters that were varied. Results for these varied parameters are given in Table S2.

| Fit | Model | Fixed parameters | | | | | |
|---|---|---|---|---|---|---|---|
| | | N1 | *K*1' ($M^{-1}$) | ΔH1 (kcal $mol^{-1}$) | N2 | *K*2' ($M^{-1}$) | ΔH2 (kcal $mol^{-1}$) |
| No FFA, 60 μM HSA | Two sets of sites | v | v | v | v | v | v |
| Others | Two sets of sites | v | 405300 | -6066 | 1.00 | v | v |

**Table S2. Fitting results from ITC experiments.**

| Fitted Parameter | Fit | 0 FFA | 2.5 FFA | 3 FFA | 4 FFA | 5 FFA |
|---|---|---|---|---|---|---|
| N1 | No FFA, 60 μM HSA | 1.000 | | | | |
| *K*1' ($M^{-1}$) | | 405300 | | | | |
| ΔH1 (kcal $mol^{-1}$) | | -6066 | | | | |
| N2 | | 1.000 | | | | |
| *K*2' ($M^{-1}$) | | 8850 | | | | |
| ΔH2 (kcal $mol^{-1}$) | | -11020 | | | | |
| N1 | Octanoate | | | 0.843 | | 0.866 |
| | Laurate | | | 0.869 | 0.244 | 4e-15 |
| | Myristate | | | 0.497 | 0.203 | 2e-14 |
| | Palmitate, 60 μM HSA | | | 0.485 | 8e-14 | 4e-18 |
| | No FFA, 25 μM HSA | 0.822 | | | | |
| | Palmitate, 25 μM HSA | | 0.788 | | 0.056 | 2e-15 |
| | Palmitoleate | | 1.000 | | 0.395 | 6e-15 |
| | Palmitelaidate | | 0.549 | | 0.318 | 1e-15 |
| | Stearate | | 0.447 | | 0.079 | 5e-20 |
| *K*2' ($M^{-1}$) | Octanoate | | | 14730 | | 13610 |
| | Laurate | | | 9310 | 10630 | 7820 |
| | Myristate | | | 11970 | 7430 | 6290 |
| | Palmitate, 60 μM HSA | | | 12170 | 7560 | 5830 |
| | No FFA, 25 μM HSA | 15120 | | | | |
| | Palmitate, 25 μM HSA | | 13200 | | 8000 | 9660 |
| | Palmitoleate | | 8523 | | 17790 | 8180 |
| | Palmitelaidate | | 6200 | | 14430 | 10830 |
| | Stearate | | 24920 | | 12420 | 13860 |
| ΔH2 (kcal $mol^{-1}$) | Octanoate | | | -7060 | | -5580 |
| | Laurate | | | -11720 | -15140 | -16050 |
| | Myristate | | | -9420 | -1.990 | -12000 |
| | Palmitate, 60 μM HSA | | | -11390 | -12740 | -12830 |
| | No FFA, 25 μM HSA | -13040 | | | | |
| | Palmitate, 25 μM HSA | | -10600 | | -20320 | -19790 |
| | Palmitoleate | | -19885 | | -14380 | -15460 |
| | Palmitelaidate | | -28520 | | -17030 | -16400 |
| | Stearate | | -12810 | | -20560 | -15890 |

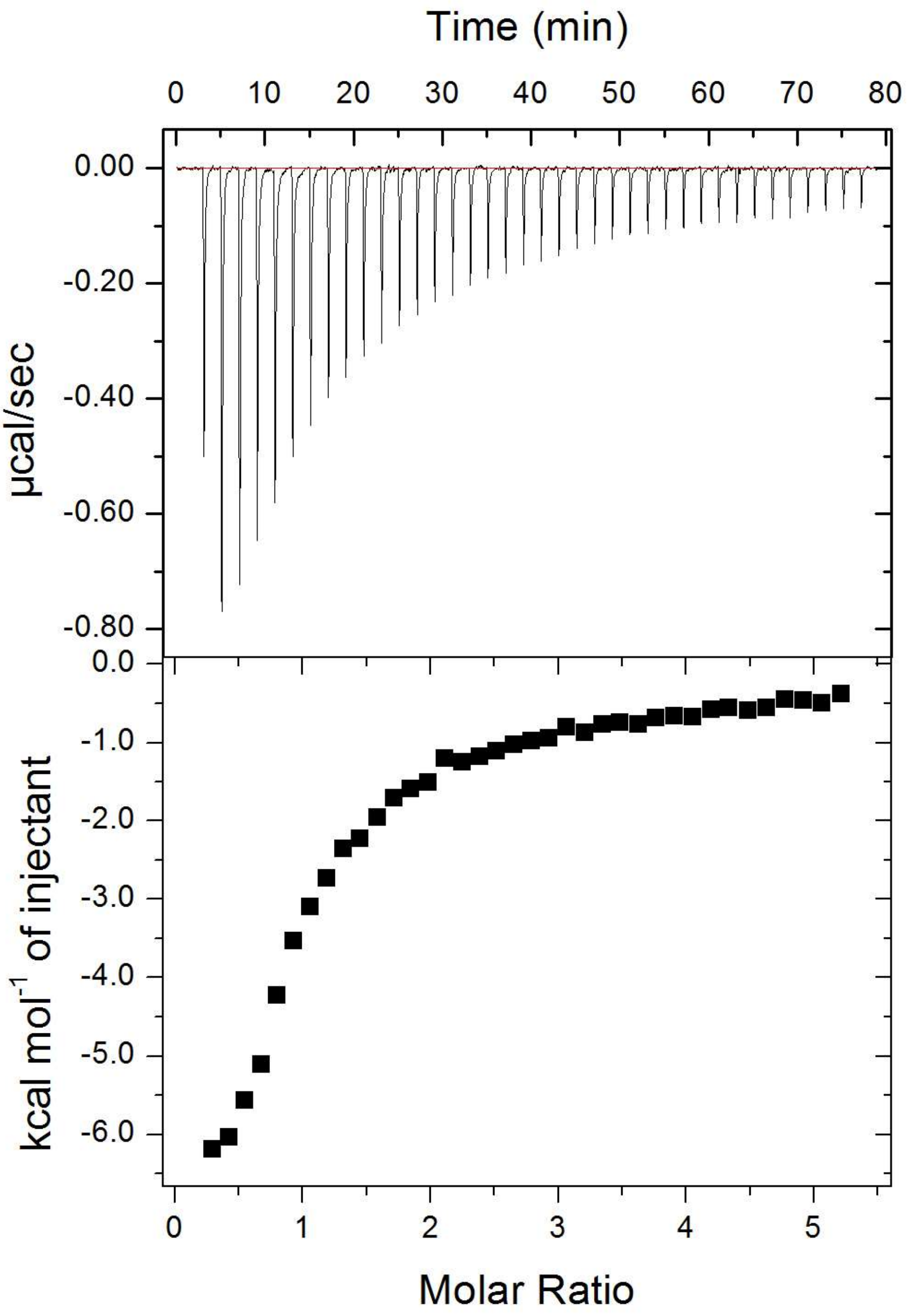


**Figure S1. ITC raw data.** Full ITC data (including raw data) for $Zn^{2+}$ binding to HSA in the absence of FFA, corresponding to data shown in Figure 2A-D.

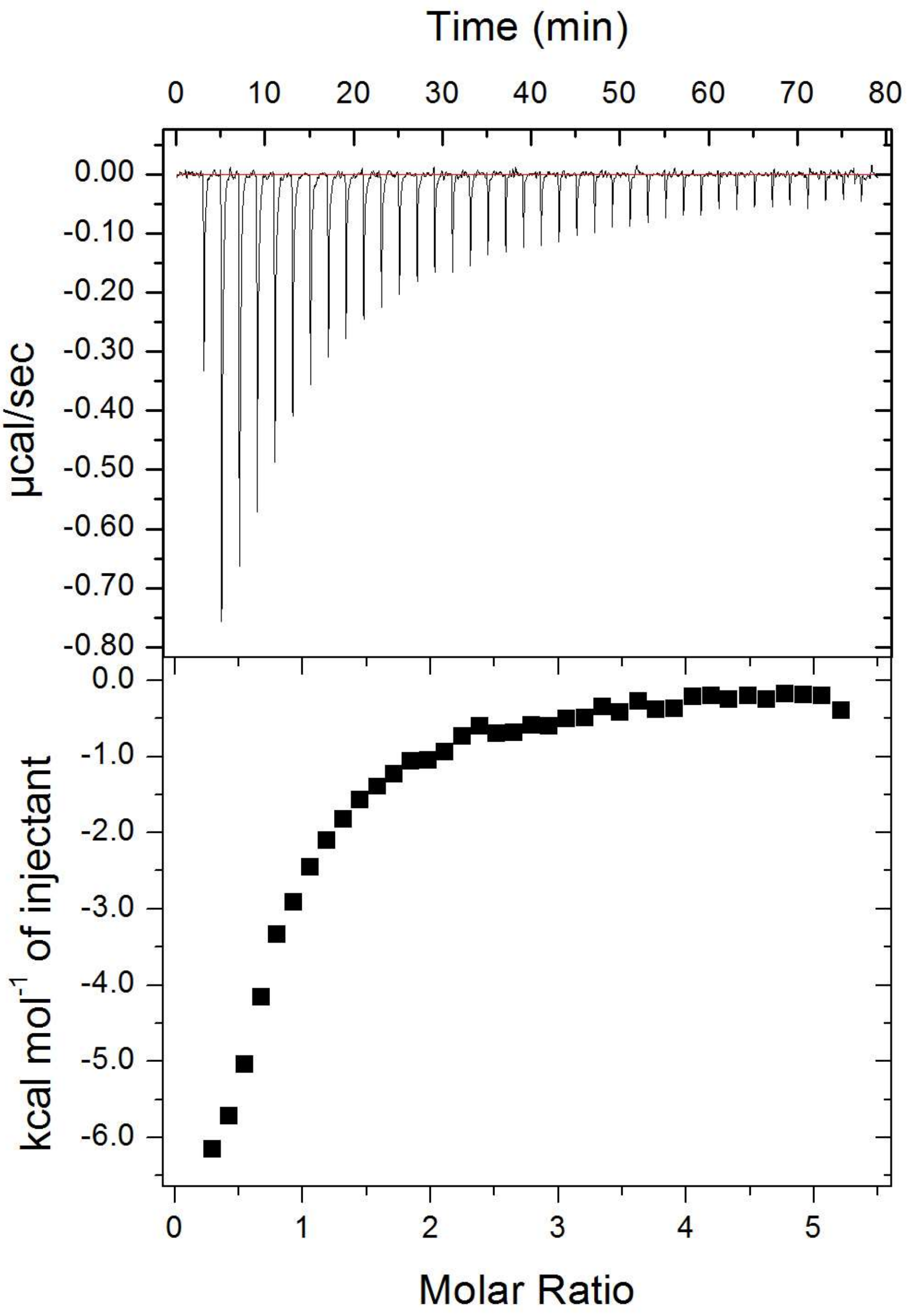


**Figure S2. ITC raw data.** Full ITC data (including raw data) for $Zn^{2+}$ binding to HSA in the presence of 3 mol. eq. of octanoate, corresponding to data shown in Figure 2A.

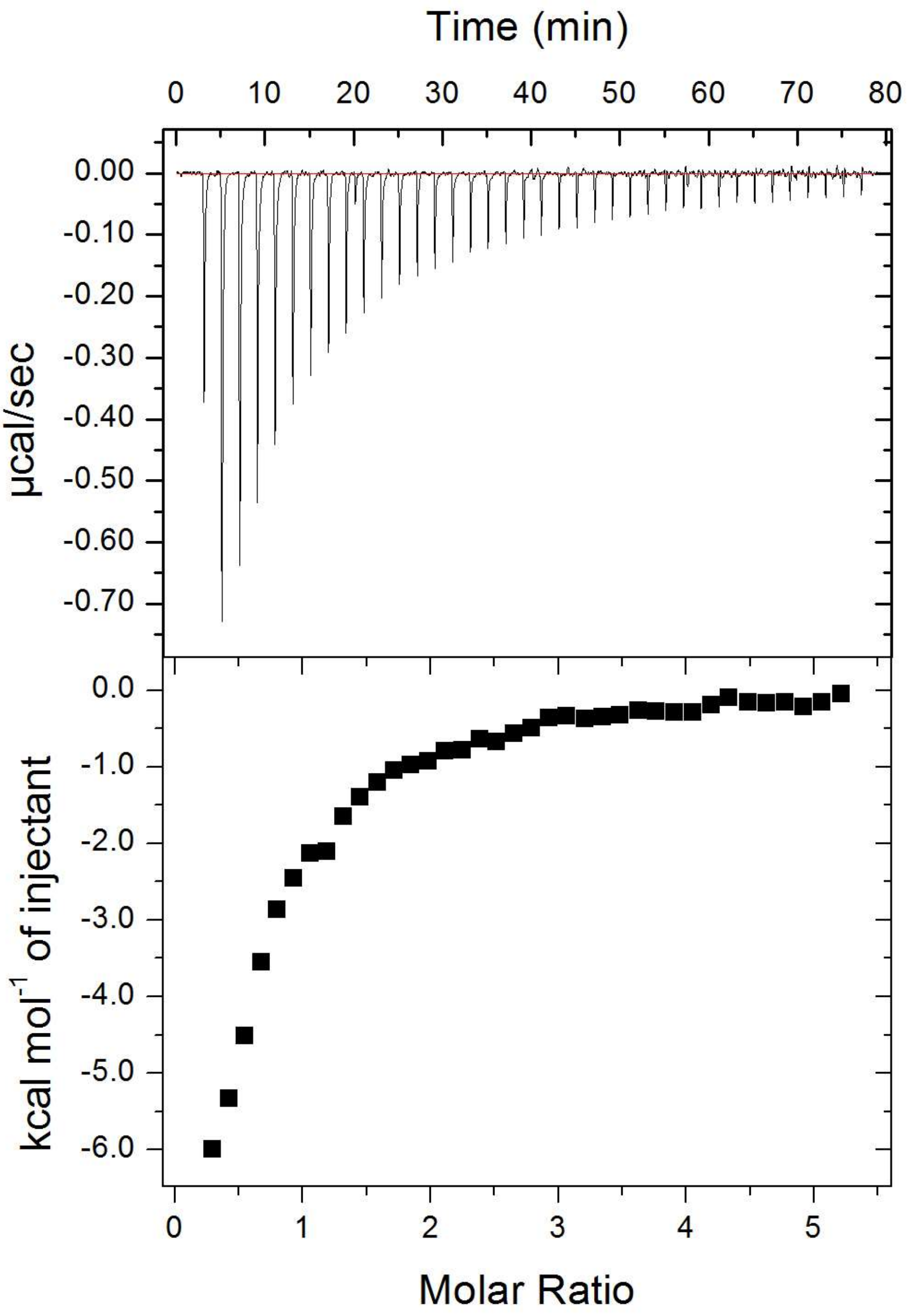


**Figure S3. ITC raw data.** Full ITC data (including raw data) for $Zn^{2+}$ binding to HSA in the presence of 5 mol. eq. of octanoate, corresponding to data shown in Figure 2A.

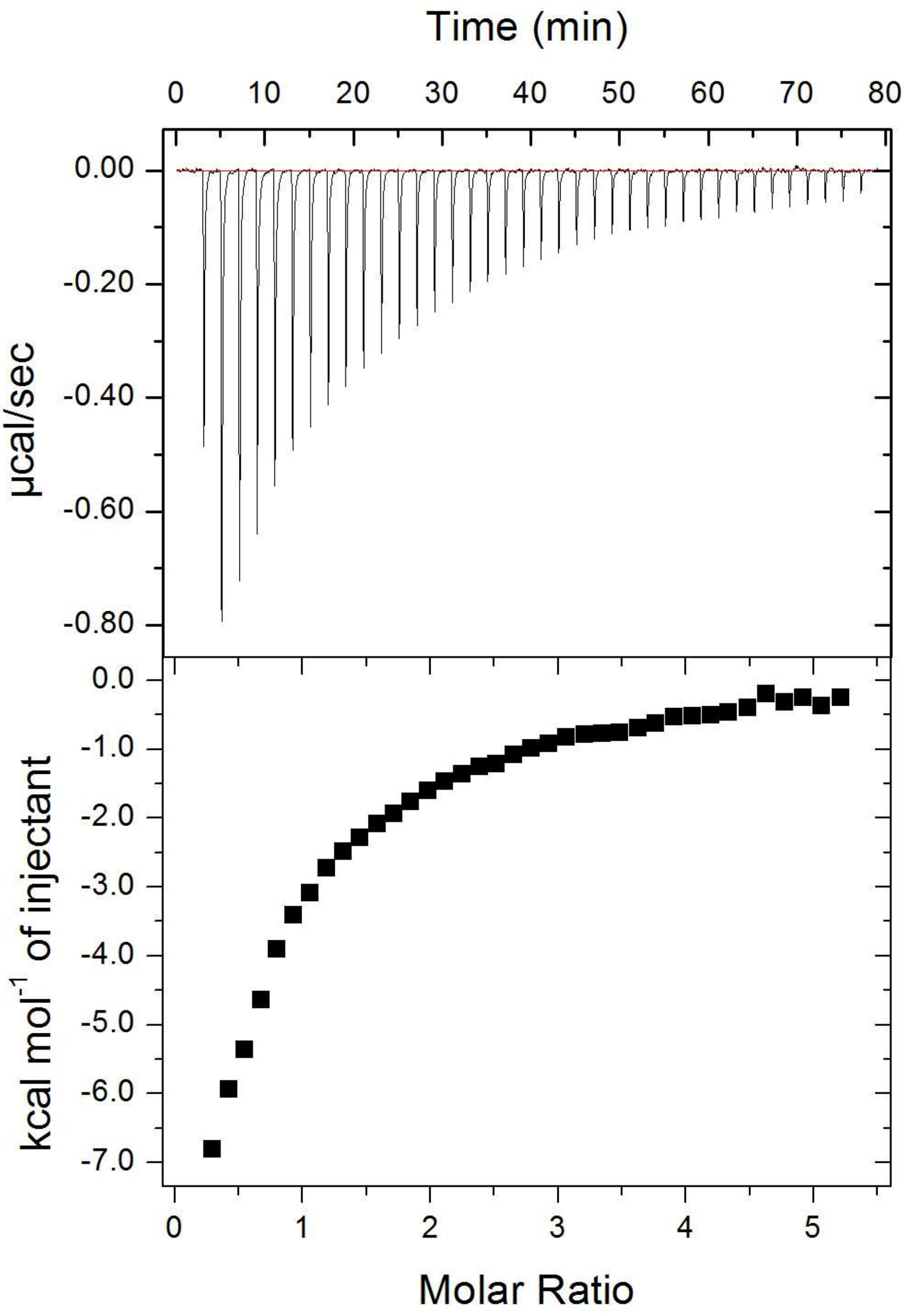


**Figure S4. ITC raw data.** Full ITC data (including raw data) for $Zn^{2+}$ binding to HSA in the presence of 3 mol. eq. of laurate, corresponding to data shown in Figure 2B.

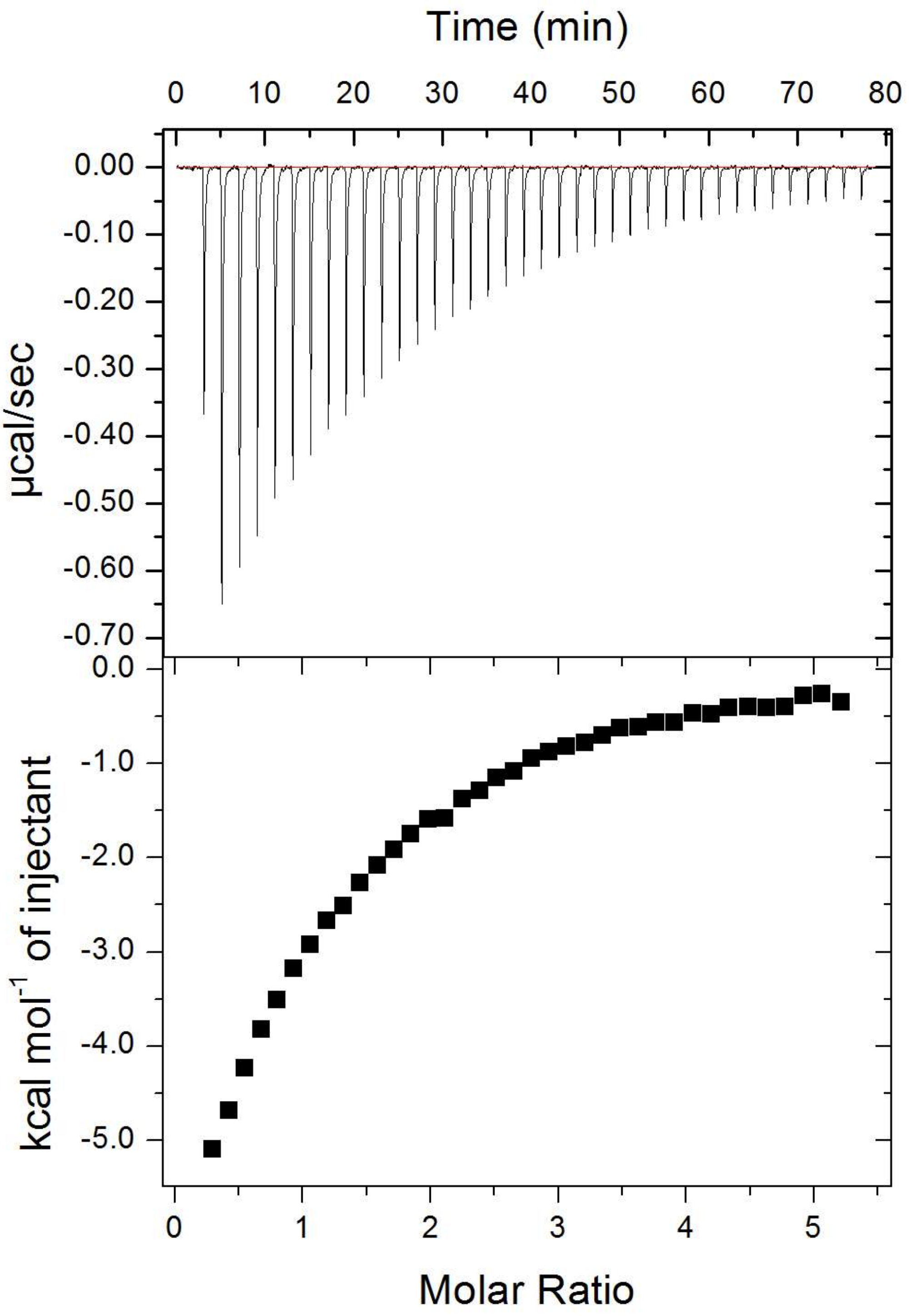


**Figure S5. ITC raw data.** Full ITC data (including raw data) for $Zn^{2+}$ binding to HSA in the presence of 4 mol. eq. of laurate, corresponding to data shown in Figure 2B.

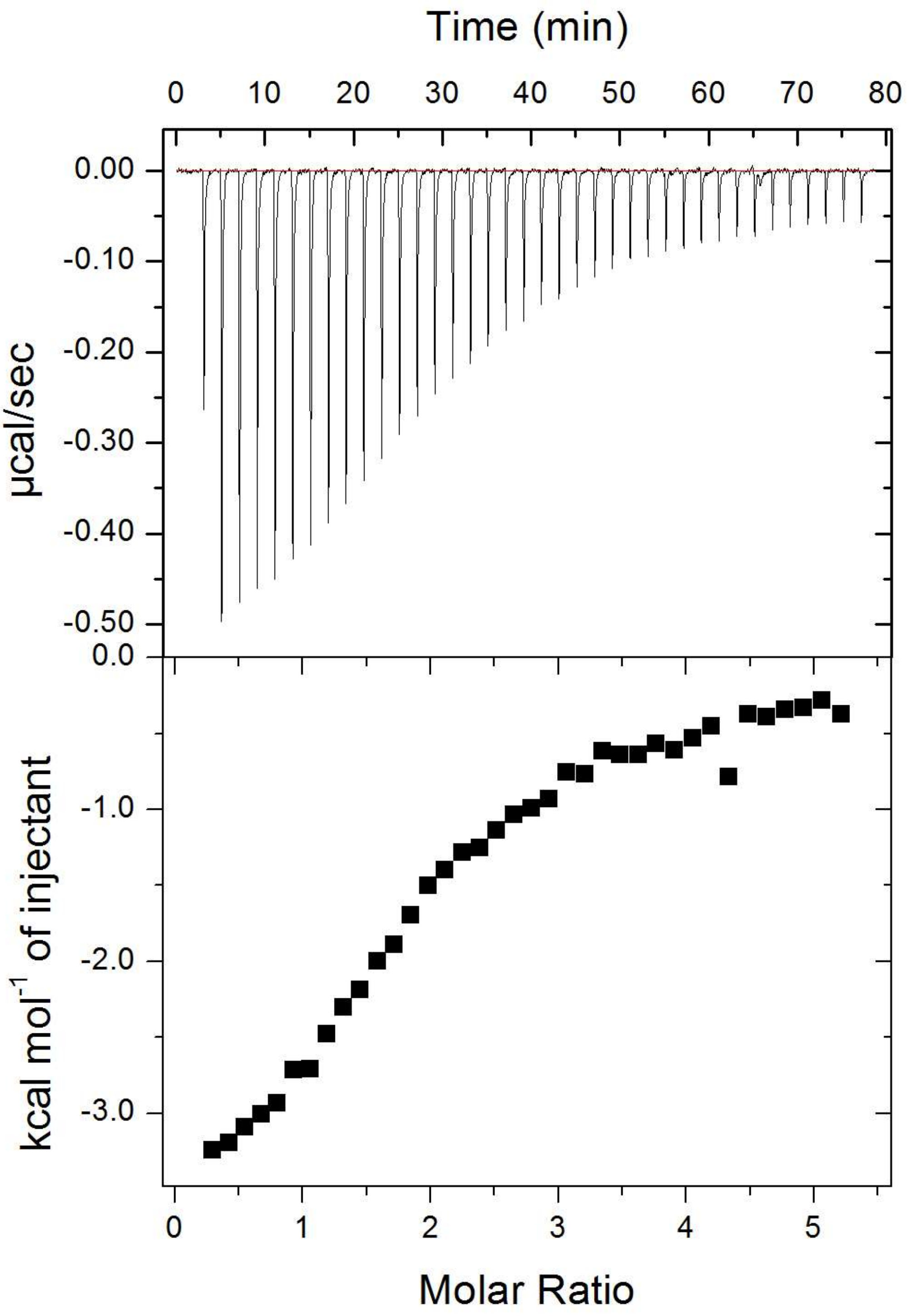


**Figure S6. ITC raw data.** Full ITC data (including raw data) for $Zn^{2+}$ binding to HSA in the presence of 5 mol. eq. of laurate, corresponding to data shown in Figure 2B.

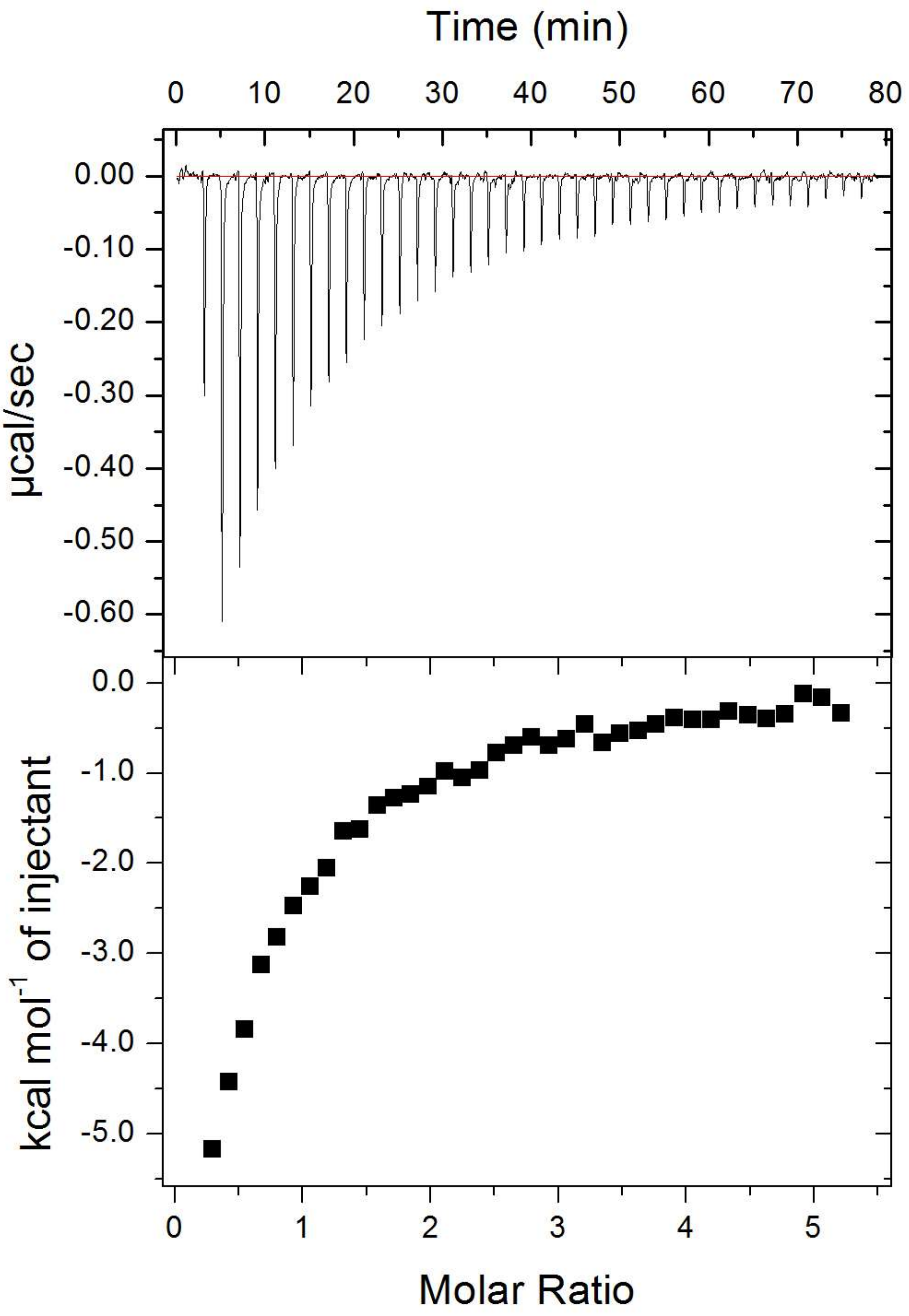


**Figure S7. ITC raw data.** Full ITC data (including raw data) for $Zn^{2+}$ binding to HSA in the presence of 3 mol. eq. of myristate, corresponding to data shown in Figure 2C.

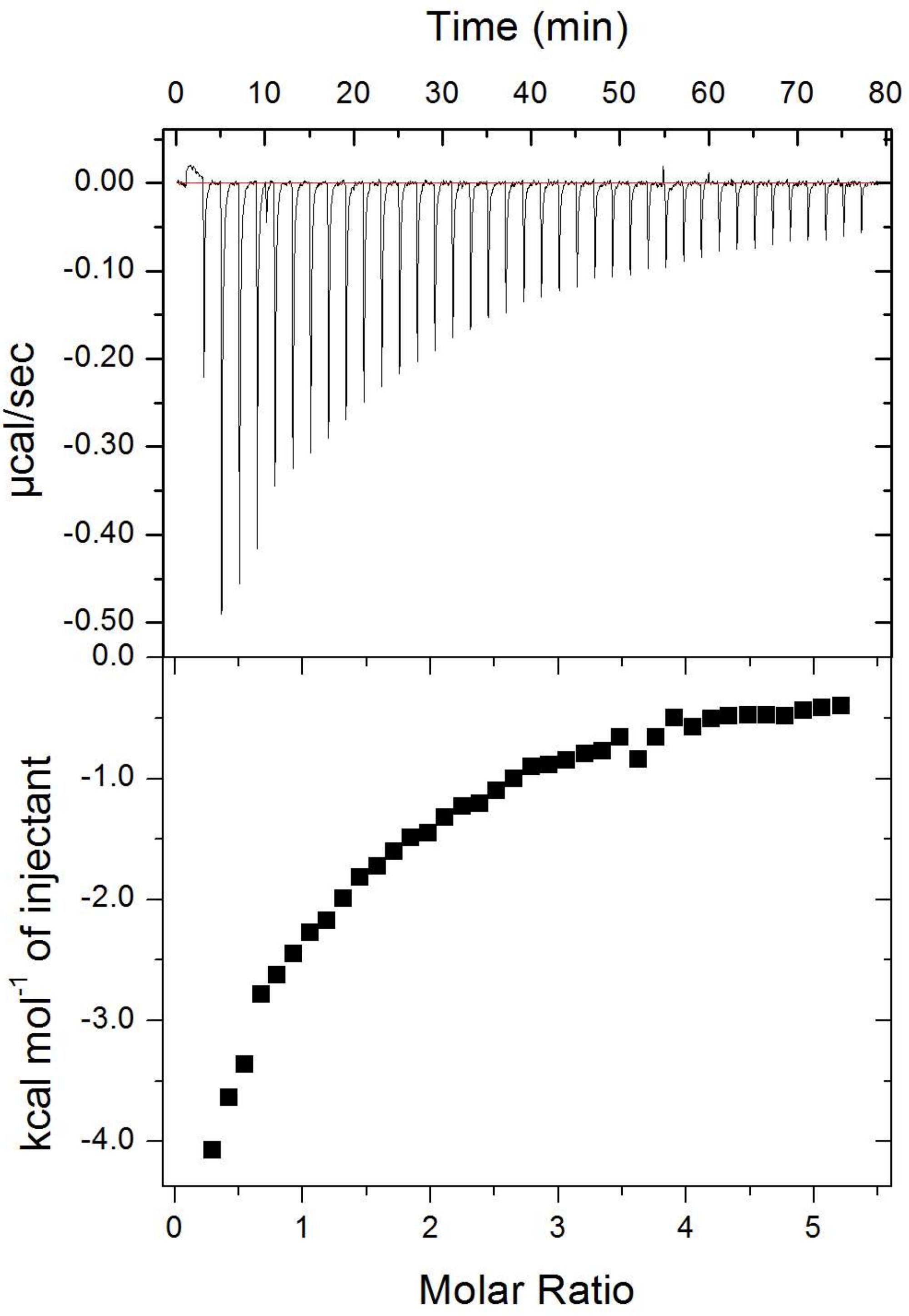


**Figure S8. ITC raw data.** Full ITC data (including raw data) for $Zn^{2+}$ binding to HSA in the presence of 4 mol. eq. of myristate, corresponding to data shown in Figure 2C.

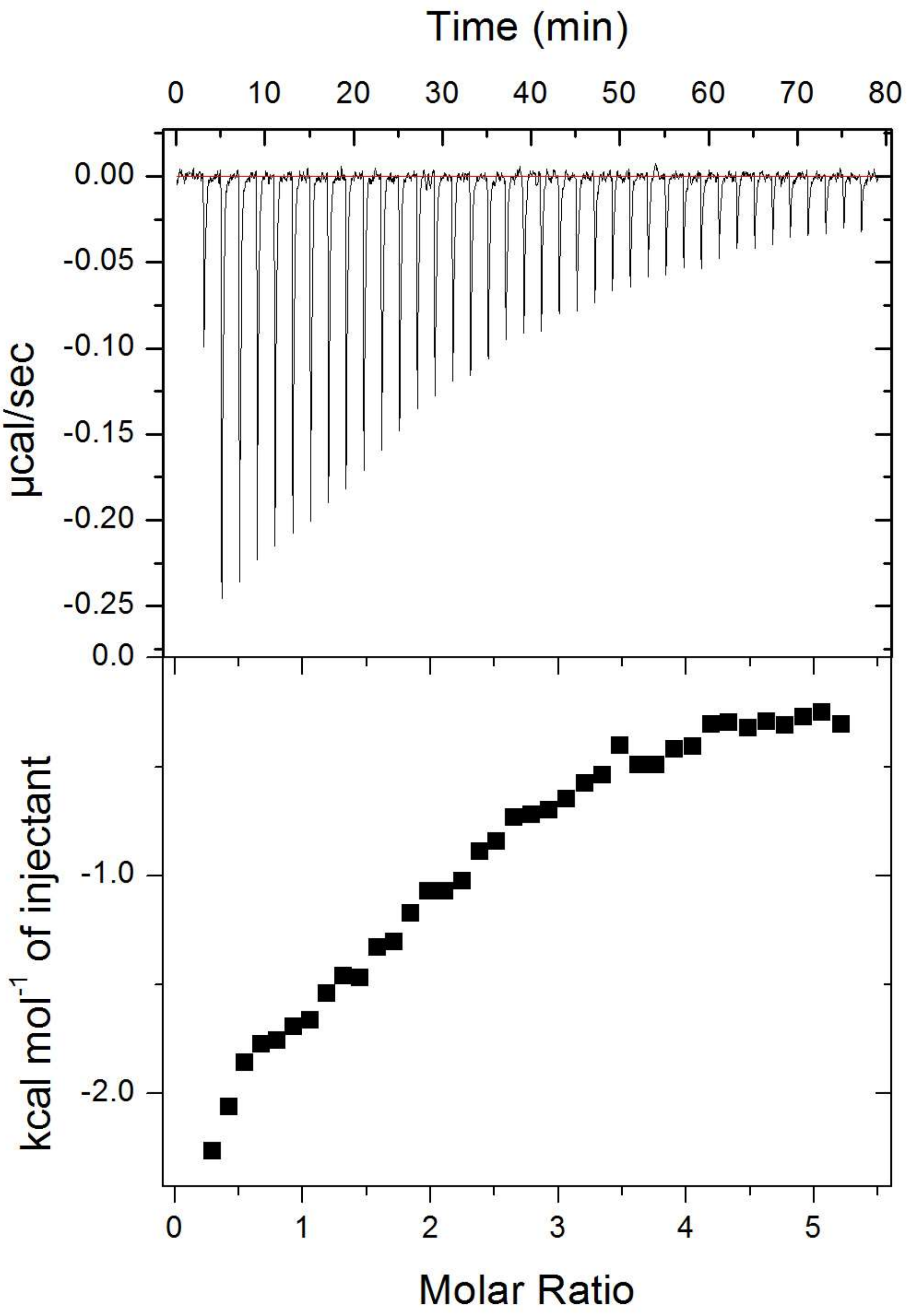


**Figure S9. ITC raw data.** Full ITC data (including raw data) for $Zn^{2+}$ binding to HSA in the presence of 5 mol. eq. of myristate, corresponding to data shown in Figure 2C.

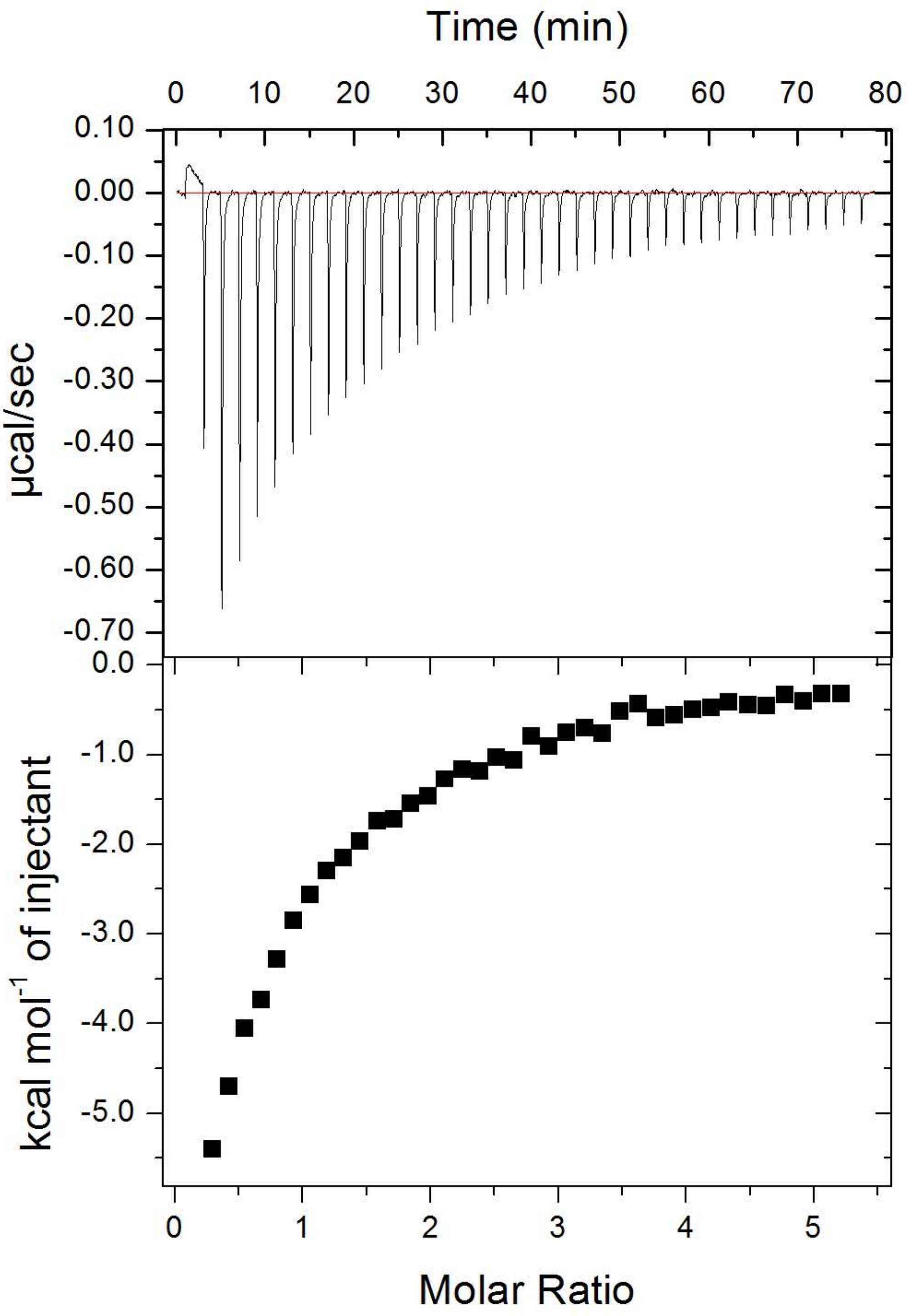


**Figure S10. ITC raw data.** Full ITC data (including raw data) for $Zn^{2+}$ binding to HSA in the presence of 3 mol. eq. of palmitate, corresponding to data shown in Figure 2D.

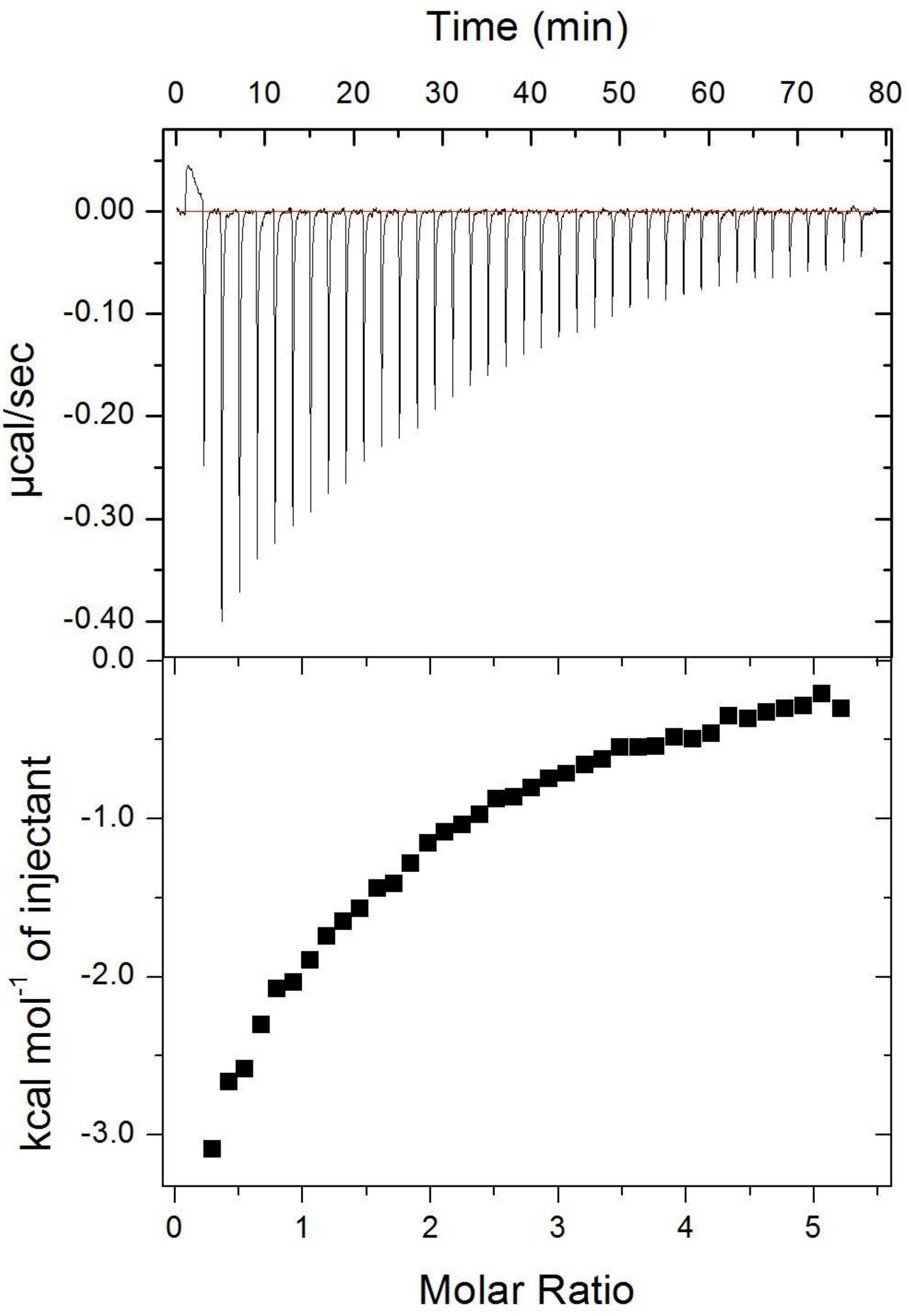


**Figure S11. ITC raw data.** Full ITC data (including raw data) for $Zn^{2+}$ binding to HSA in the presence of 4 mol. eq. of palmitate, corresponding to data shown in Figure 2D.

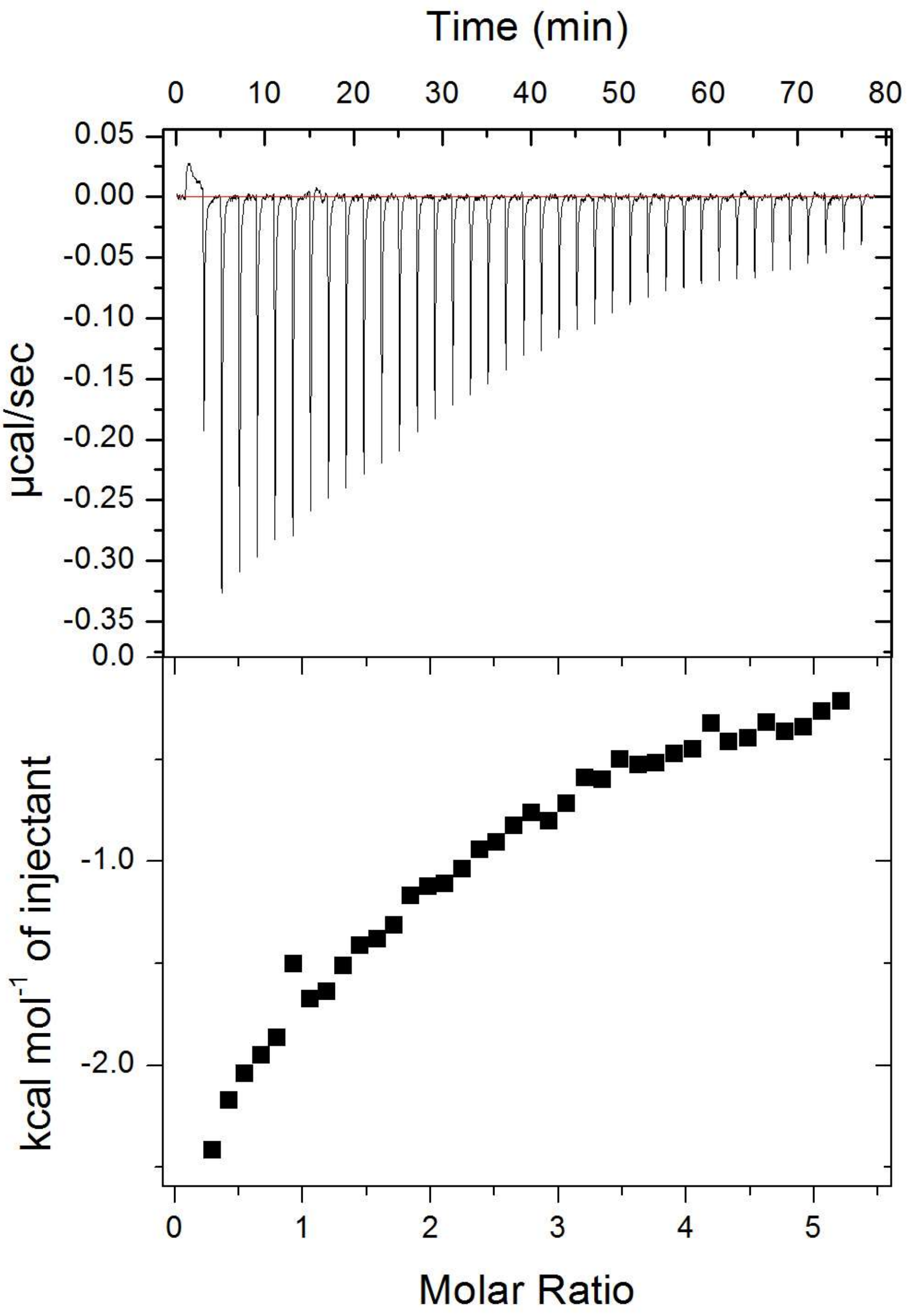


**Figure S12. ITC raw data.** Full ITC data (including raw data) for $Zn^{2+}$ binding to HSA in the presence of 5 mol. eq. of palmitate, corresponding to data shown in Figure 2D.

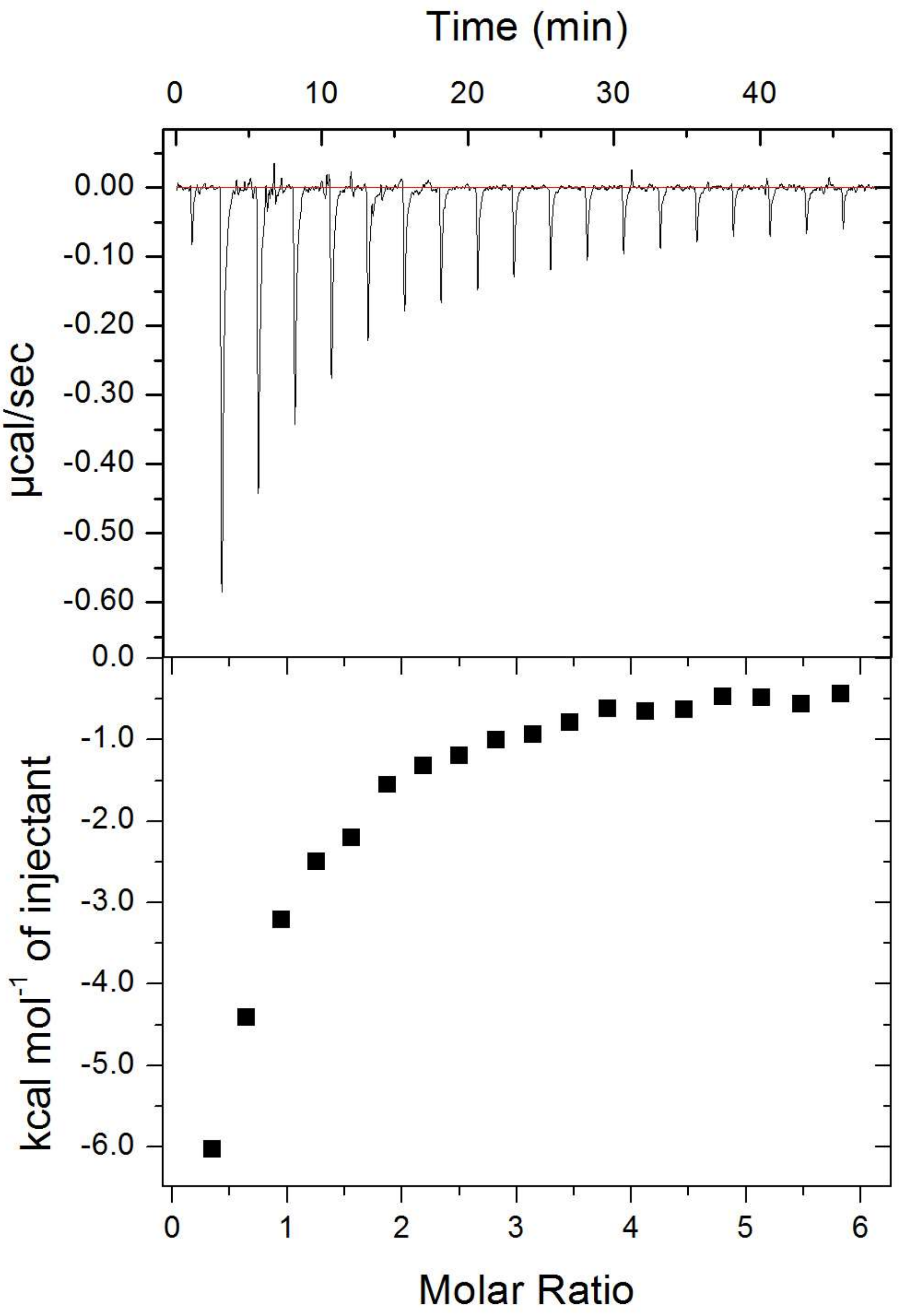


**Figure S13. ITC raw data.** Full ITC data (including raw data) for $Zn^{2+}$ binding to HSA in the absence of FFA, corresponding to data shown in Figures 2E-H.

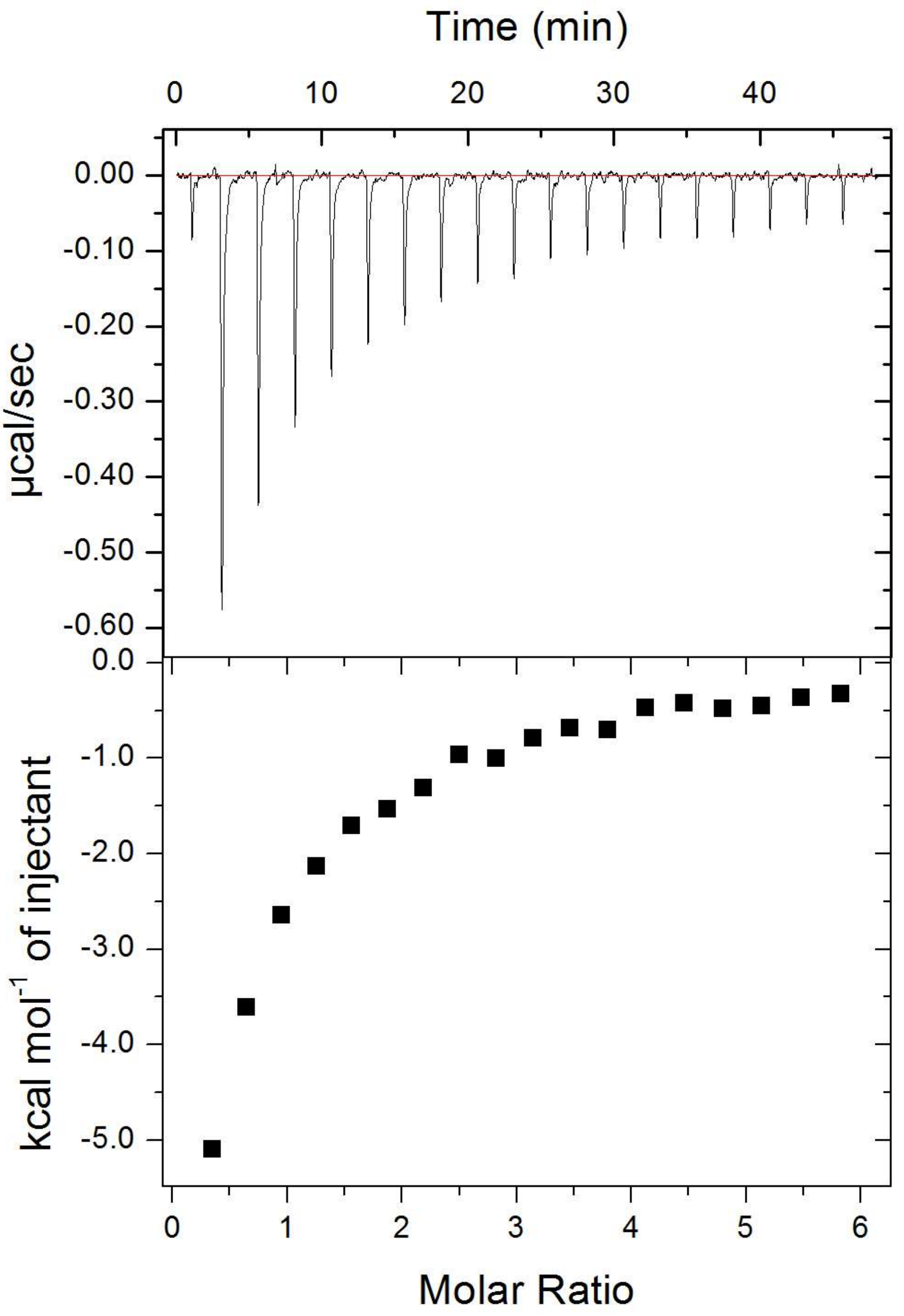


**Figure S14. ITC raw data.** Full ITC data (including raw data) for $Zn^{2+}$ binding to HSA in the presence of 2.5 mol. eq. of palmitate, corresponding to data shown in Figure 2E.

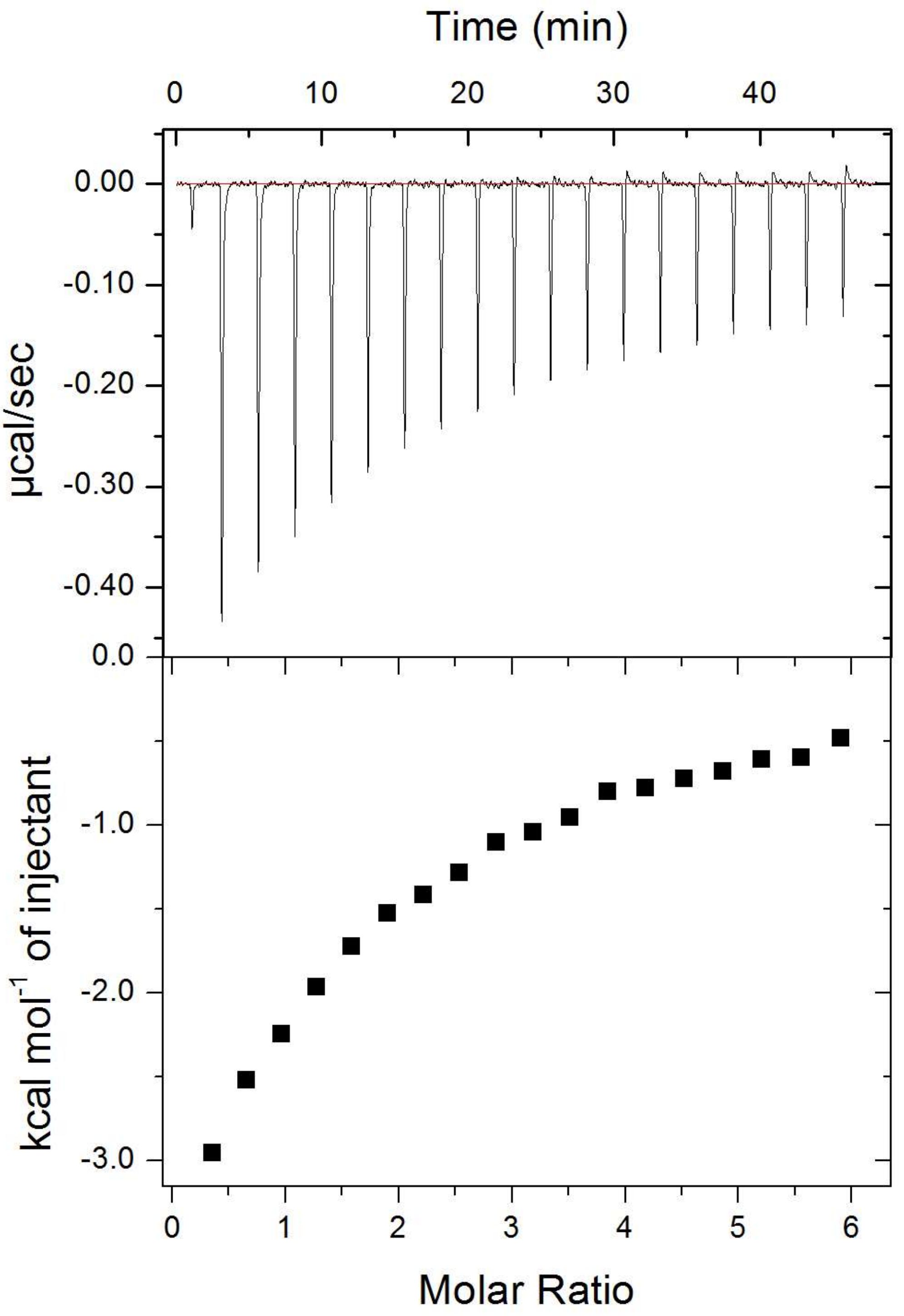


**Figure S15. ITC raw data.** Full ITC data (including raw data) for $Zn^{2+}$ binding to HSA in the presence of 4 mol. eq. of palmitate, corresponding to data shown in Figure 2E.

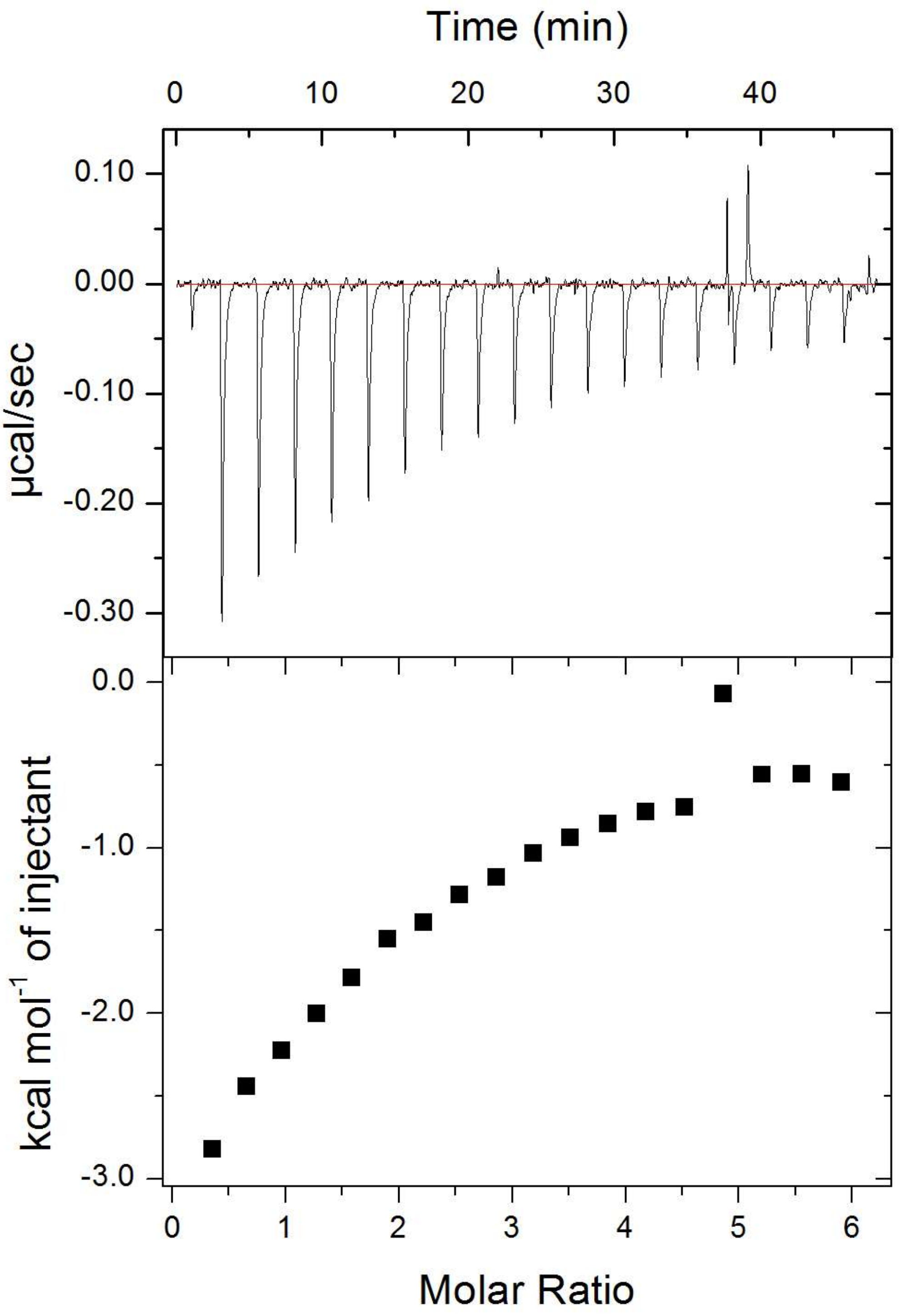


**Figure S16. ITC raw data.** Full ITC data (including raw data) for $Zn^{2+}$ binding to HSA in the presence of 5 mol. eq. of palmitate, corresponding to data shown in Figure 2E.

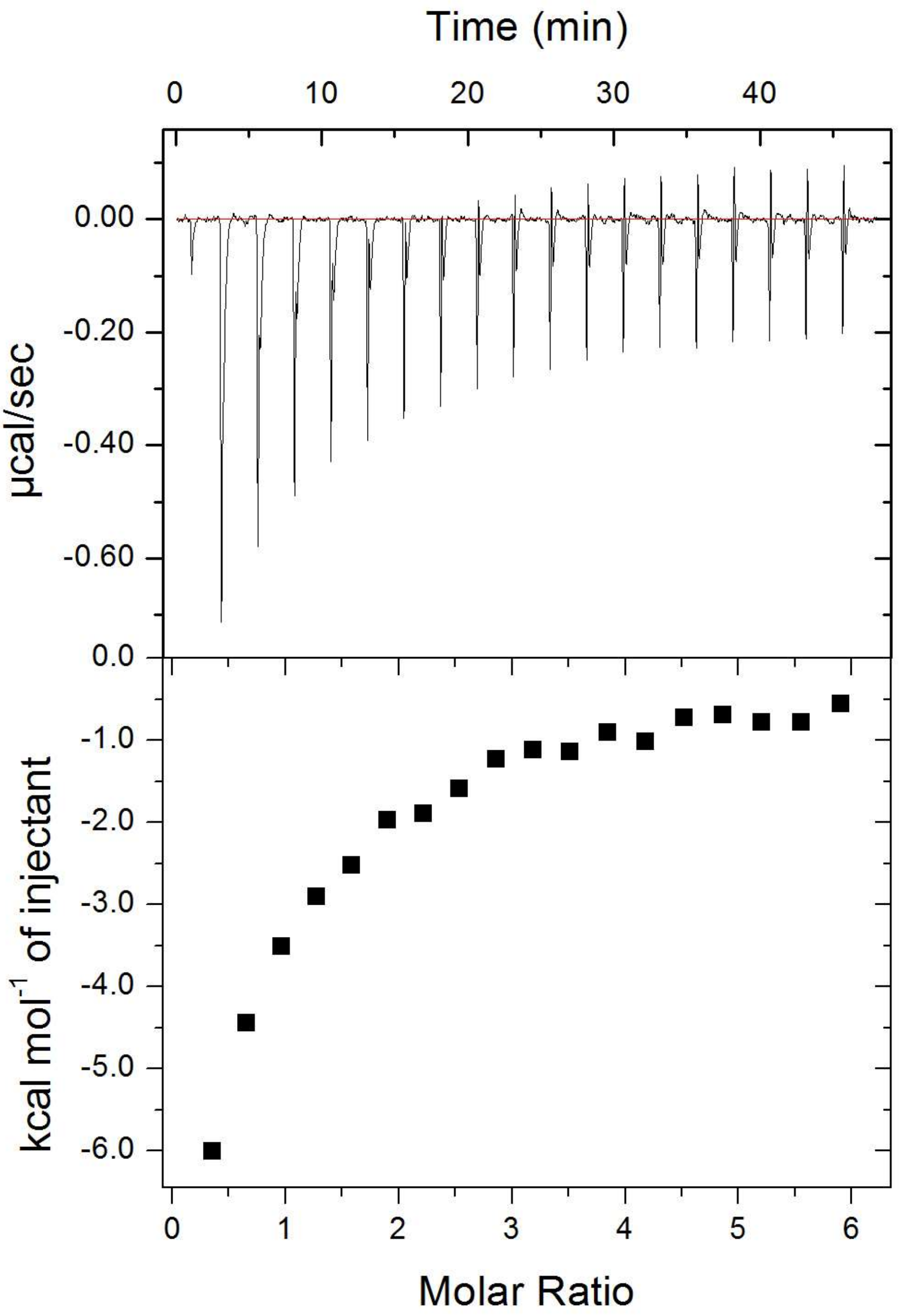


**Figure S17. ITC raw data.** Full ITC data (including raw data) for $Zn^{2+}$ binding to HSA in the presence of 2.5 mol. eq. of palmitoleate, corresponding to data shown in Figure 2F.

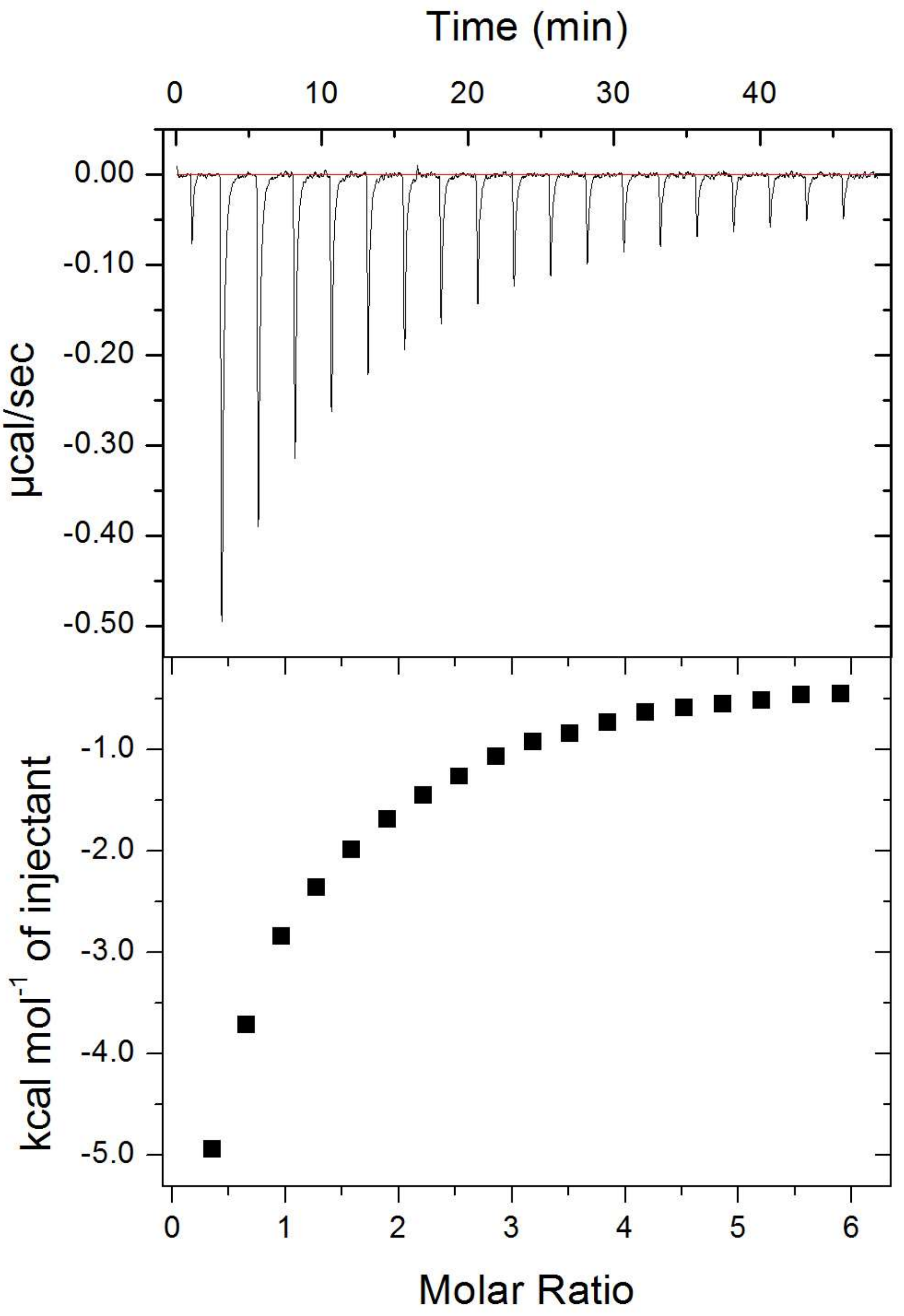


**Figure S18. ITC raw data.** Full ITC data (including raw data) for $Zn^{2+}$ binding to HSA in the presence of 4 mol. eq. of palmitoleate, corresponding to data shown in Figure 2F.

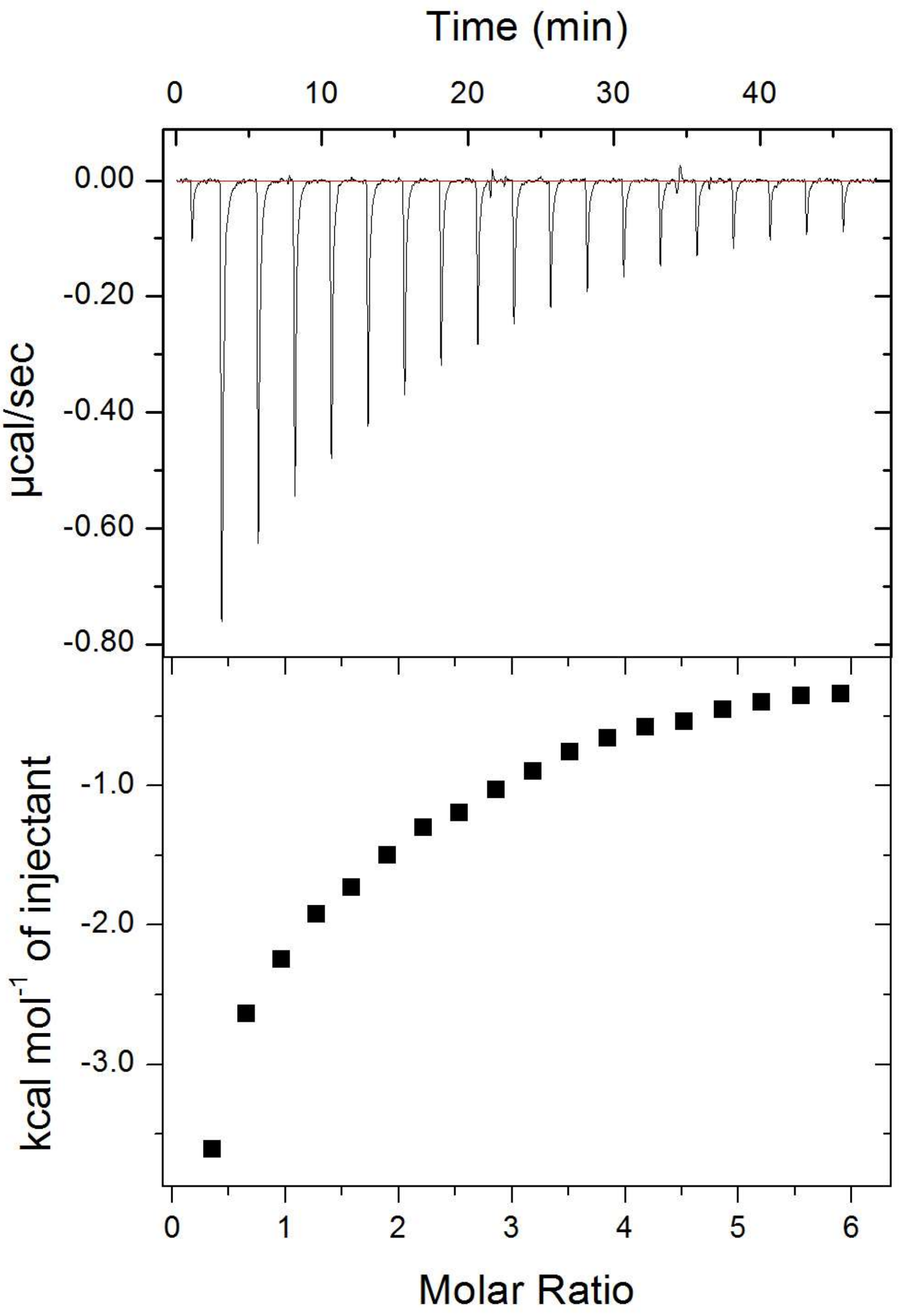


**Figure S19. ITC raw data.** Full ITC data (including raw data) for $Zn^{2+}$ binding to HSA in the presence of 5 mol. eq. of palmitoleate, corresponding to data shown in Figure 2F.

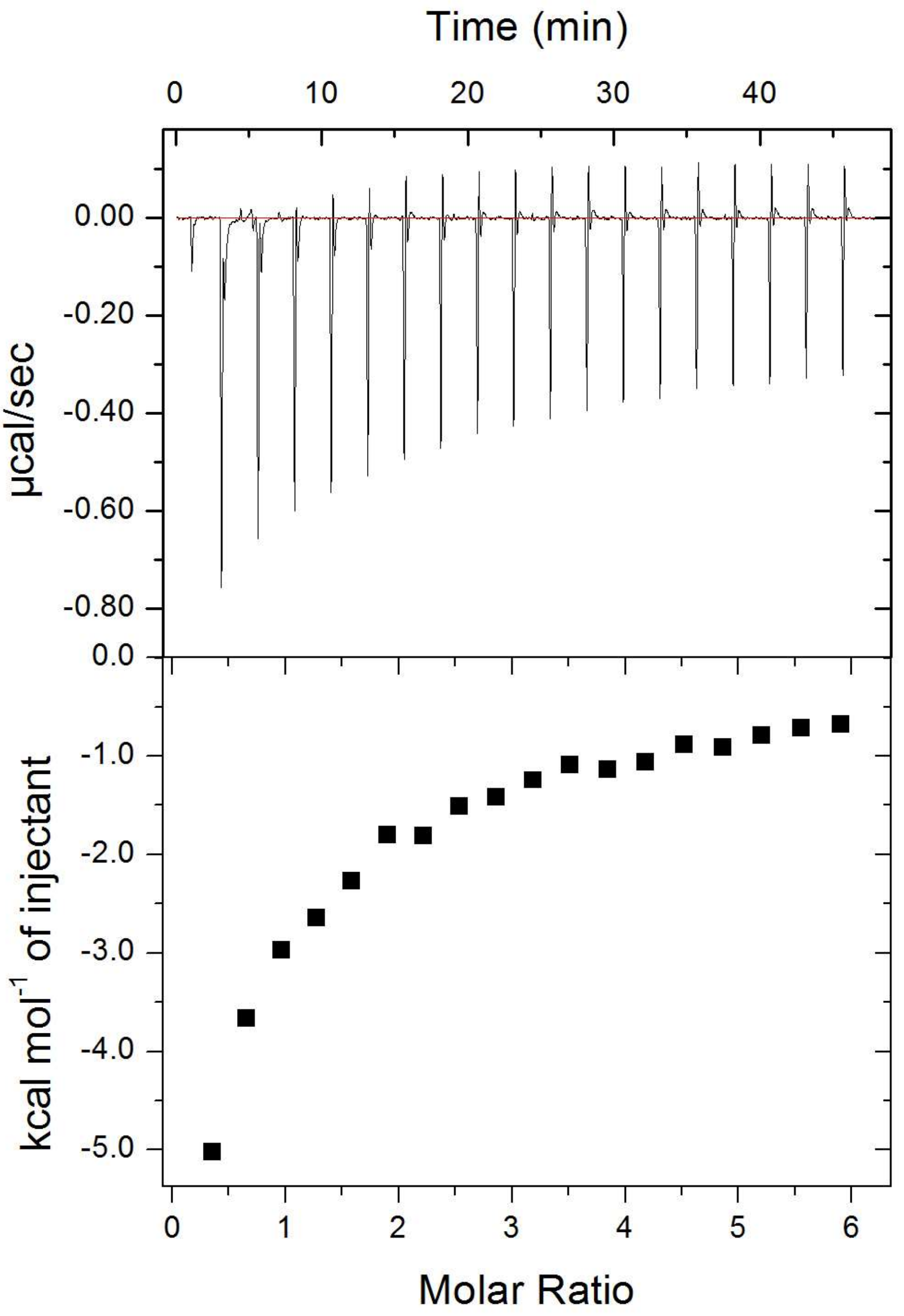


**Figure S20. ITC raw data.** Full ITC data (including raw data) for $Zn^{2+}$ binding to HSA in the presence of 2.5 mol. eq. of palmitelaidate, corresponding to data shown in Figure 2G.

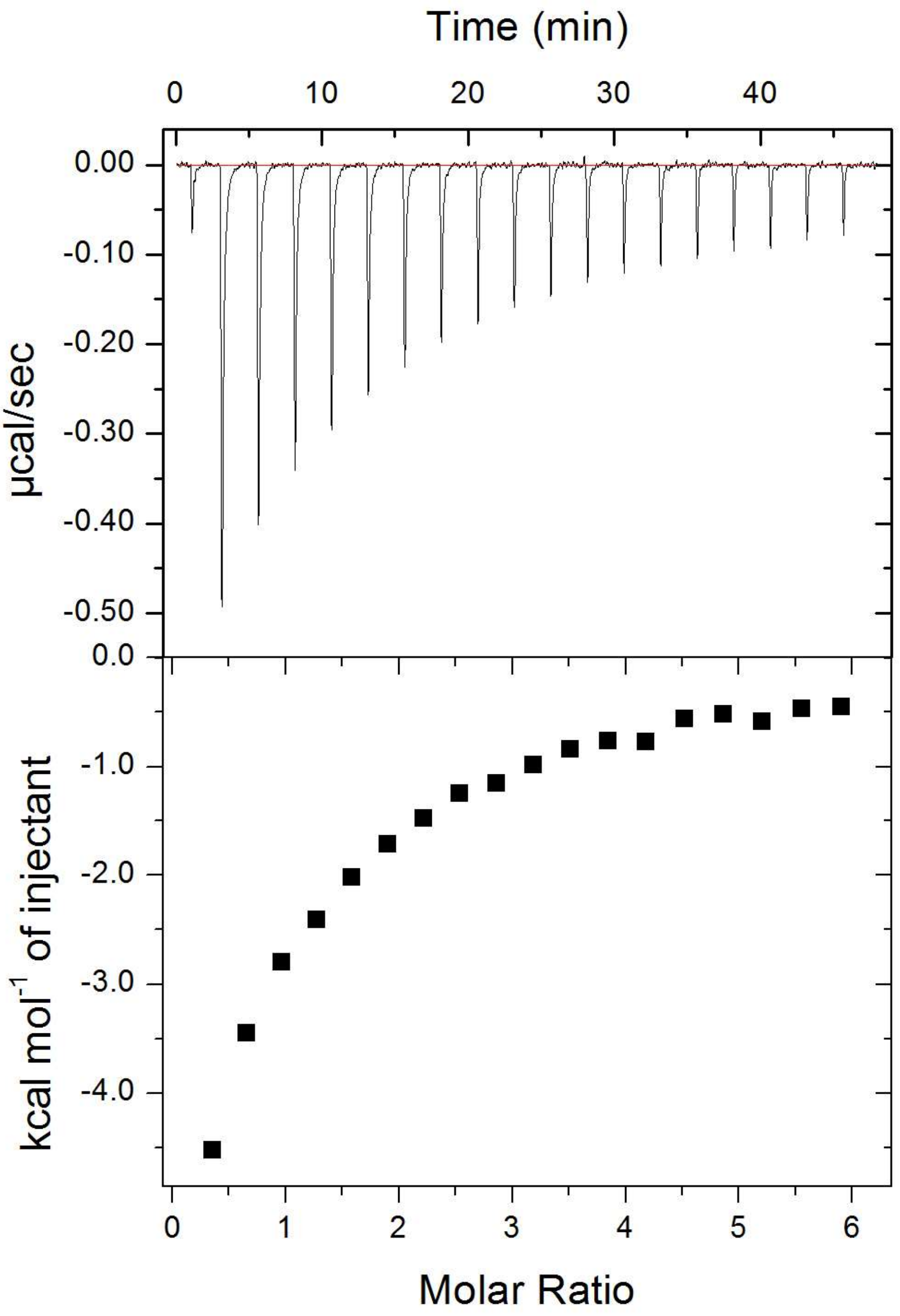


**Figure S21. ITC raw data.** Full ITC data (including raw data) for $Zn^{2+}$ binding to HSA in the presence of 4mol. eq. of palmitelaidate, corresponding to data shown in Figure 2G.

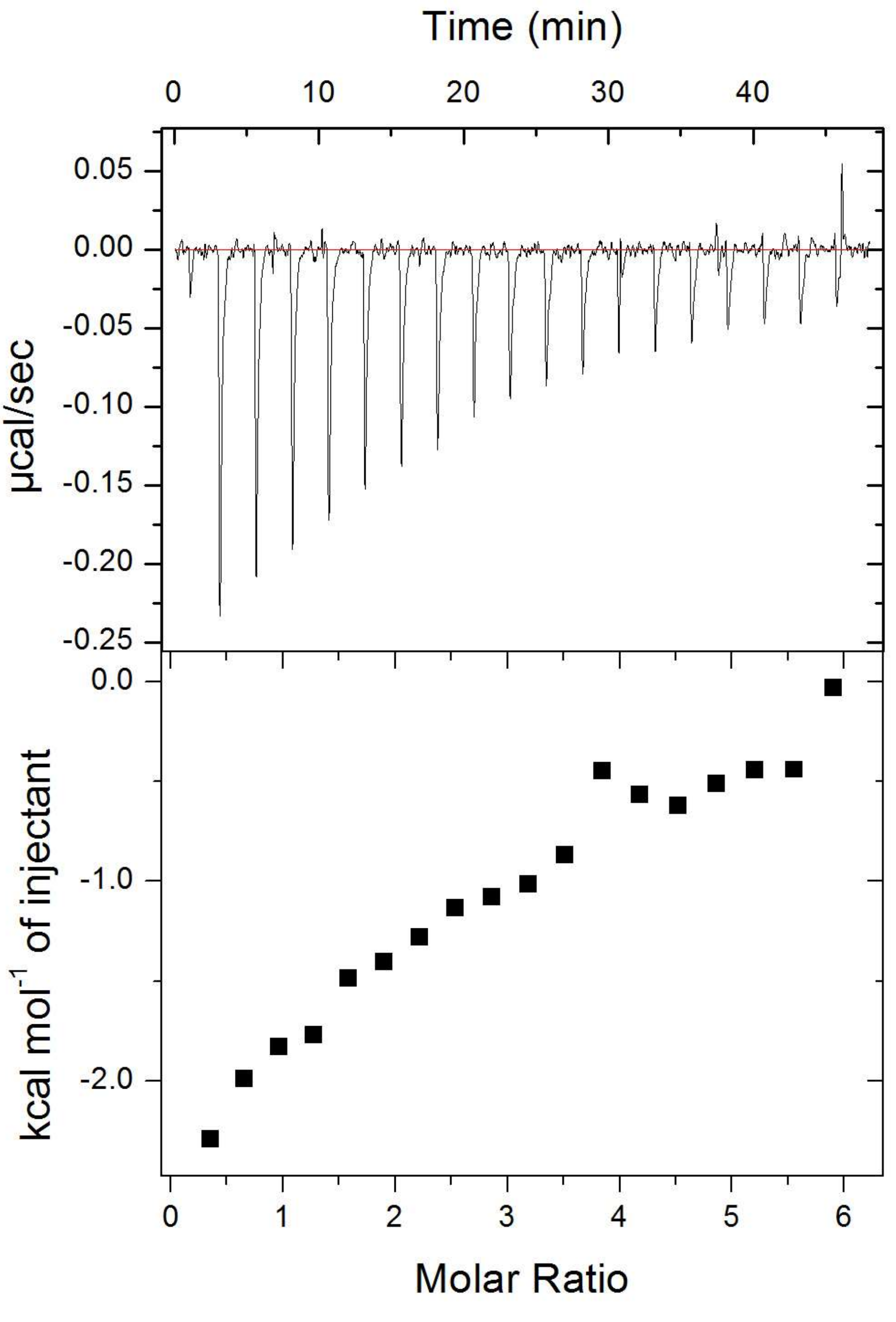


**Figure S22. ITC raw data.** Full ITC data (including raw data) for $Zn^{2+}$ binding to HSA in the presence of 5 mol. eq. of palmitelaidate, corresponding to data shown in Figure 2G.

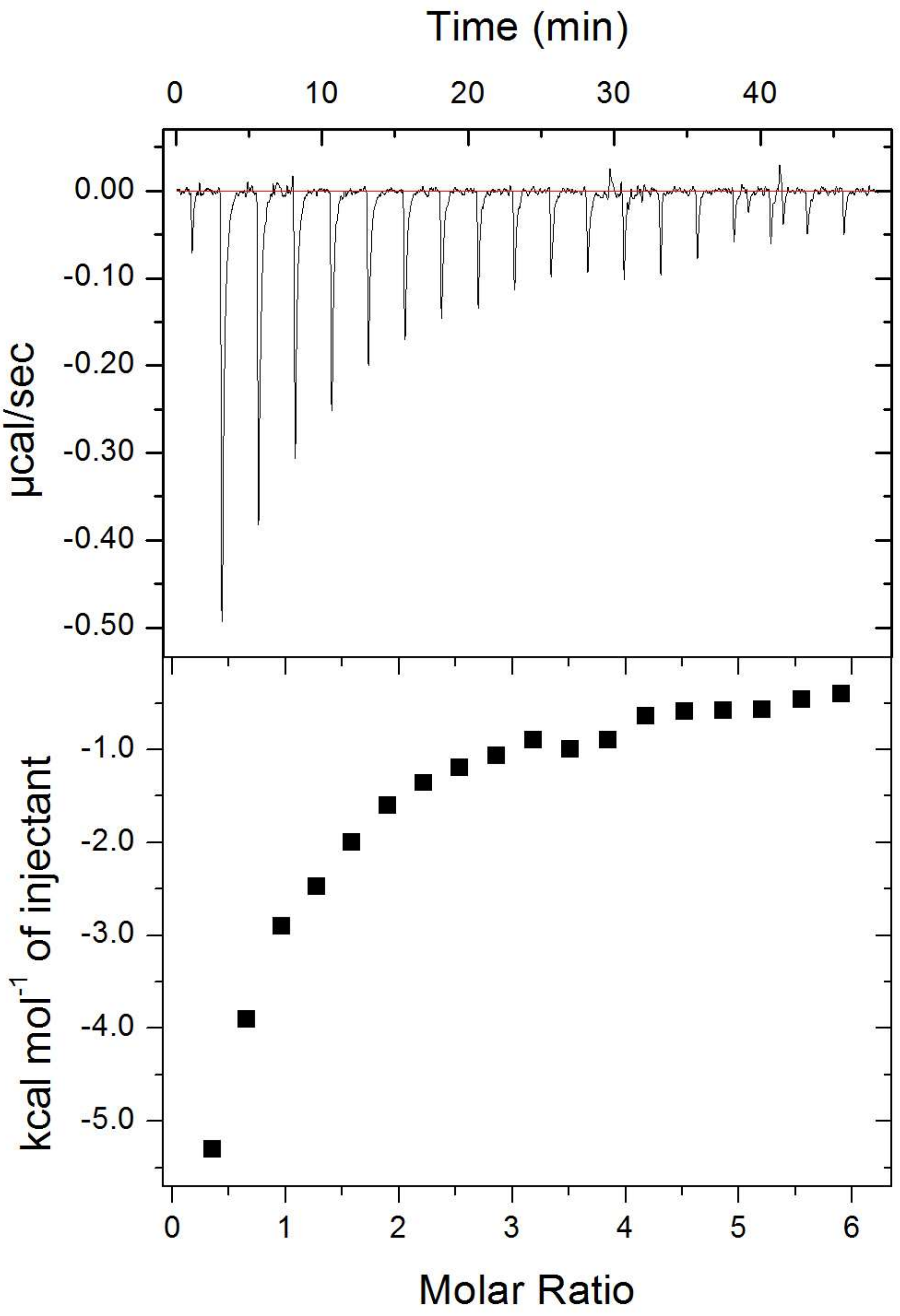


**Figure S23. ITC raw data.** Full ITC data (including raw data) for $Zn^{2+}$ binding to HSA in the presence of 2.5 mol. eq. of stearate, corresponding to data shown in Figure 2H.

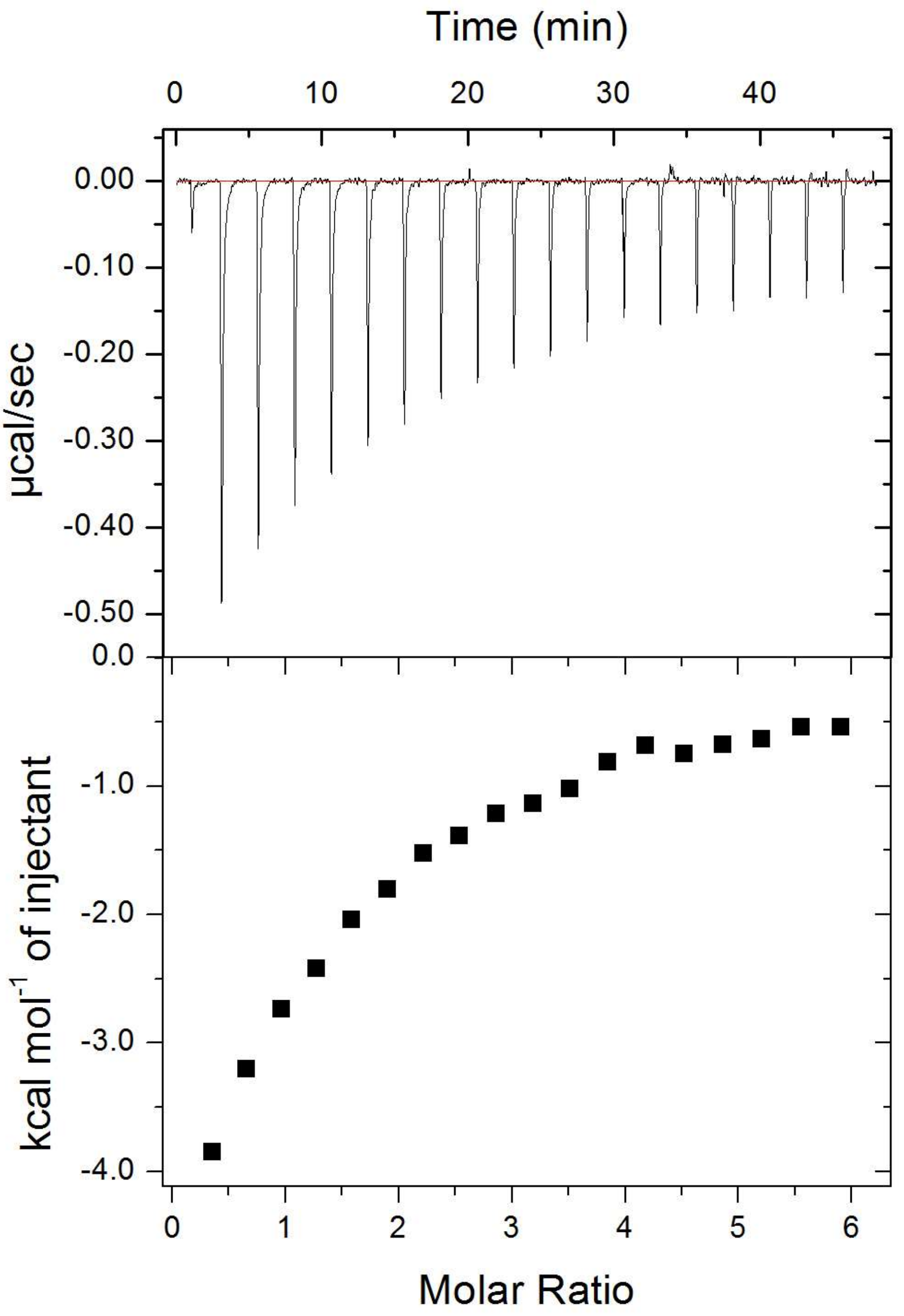


**Figure S24. ITC raw data.** Full ITC data (including raw data) for $Zn^{2+}$ binding to HSA in the presence of 4 mol. eq. of stearate, corresponding to data shown in Figure 2H.

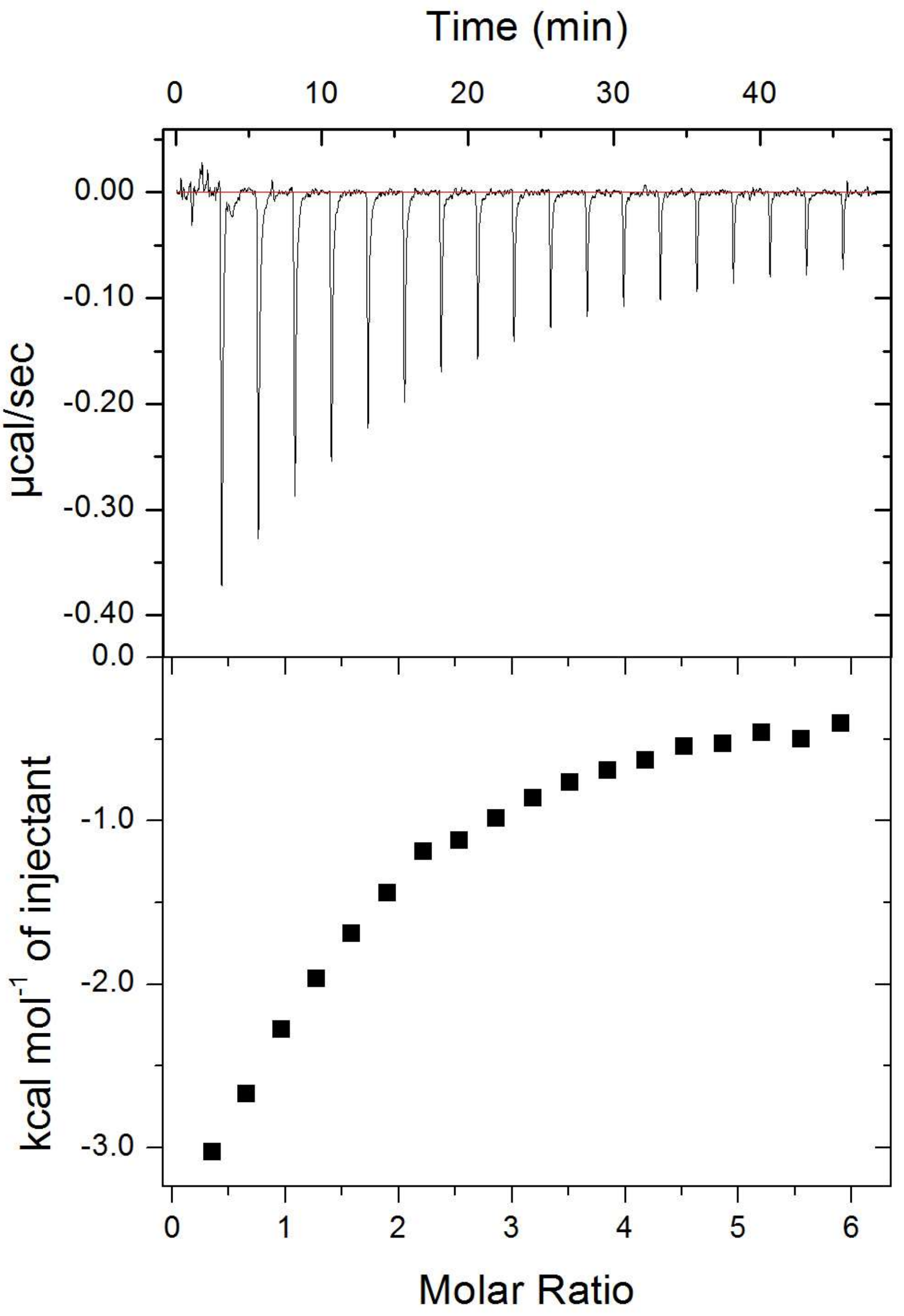


**Figure S25. ITC raw data.** Full ITC data (including raw data) for $Zn^{2+}$ binding to HSA in the presence of 5 mol. eq. of stearate, corresponding to data shown in Figure 2H.

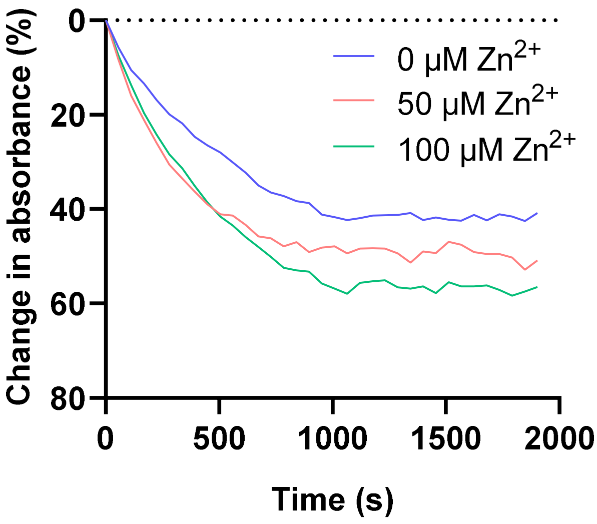


**Figure S26. Representative raw data from platelet aggregation assays.** Washed platelets, no myristate, 0-100 μM $Zn^{2+}$.

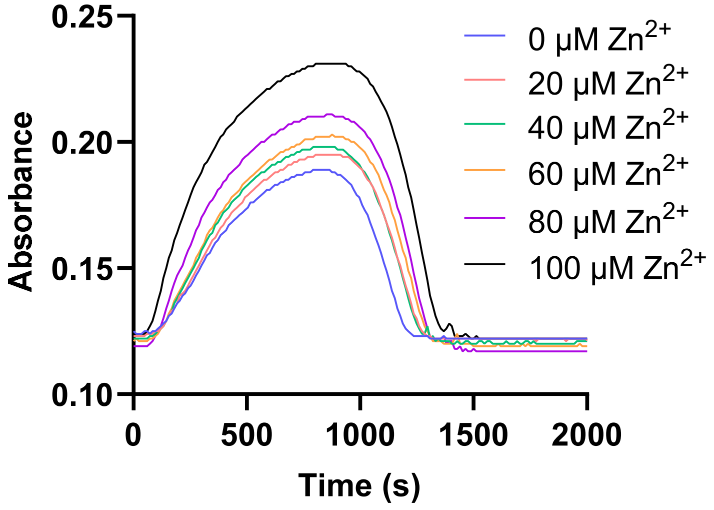


**Figure S27. Representative raw data from turbidimetric fibrin clotting and lysis assays.** Purified system, no myristate, 0-100 μM $Zn^{2+}$.

**Table S3. Demographic information and the results from plasma analysis for the T2DM and control subjects.** All values are presented as mean ± standard deviation. The *P*-values were calculated by Student's *t*-test.

| | Controls (n=18) | Patients with T2DM (n=54) | P-values |
|---|---|---|---|
| Age (years) | 57.1 ± 8.9 | 60.9 ± 7.6 | 0.0814 |
| Sex (% female) | 56 | 13 | 0.0001 |
| Weight (kg) | 70.2 ± 12.9 | 96.3 ± 17.8 | <0.0001 |
| BMI | 25.0 ± 3.2 | 32.6 ± 5.3 | <0.0001 |
| Number of individuals who smoke | 1 | 10 | - |
| Number of individuals who have had microvascular events | - | 22 | - |
| Number of individuals who have had macrovascular events | - | 31 | - |
| Numbers of individuals with familial history of autoimmunity | 2 | 7 | - |
| Numbers of individuals with familial history of Huntington's disease | 3 | 30 | - |
| Diabetes duration (months) | - | 139 ± 78 | - |
| Fasting glucose (mM) | 4.8 ± 0.5 | 10.3 ± 4.6 | <0.0001 |
| HbA1c (mmol/mol) | 37.6 ± 4.3 | 72.4 ± 22.8 | <0.0001 |
| Triglyceride (mM) | 0.95 ± 0.28 | 2.1 ± 2.0 | 0.0313 |
| Cholesterol (mM) | 5.3 ± 0.7 | 3.8 ± 0.8 | <0.0001 |
| LDL (mM) | 3.0 ± 0.7 | 1.9 ± 0.5 | <0.0001 |
| HDL (mM) | 1.9 ± 0.5 | 1.1 ± 0.3 | <0.0001 |
| Cholesterol/HDL ratio | 3.1 ± 0.9 | 3.7 ± 0.9 | 0.0198 |
| Platelet count | - | 251 ± 53 | - |
| Fibrinogen | 2.7 ± 0.3* | 2.7 ± 0.5 | 0.9005 |

* data only available for 3 samples

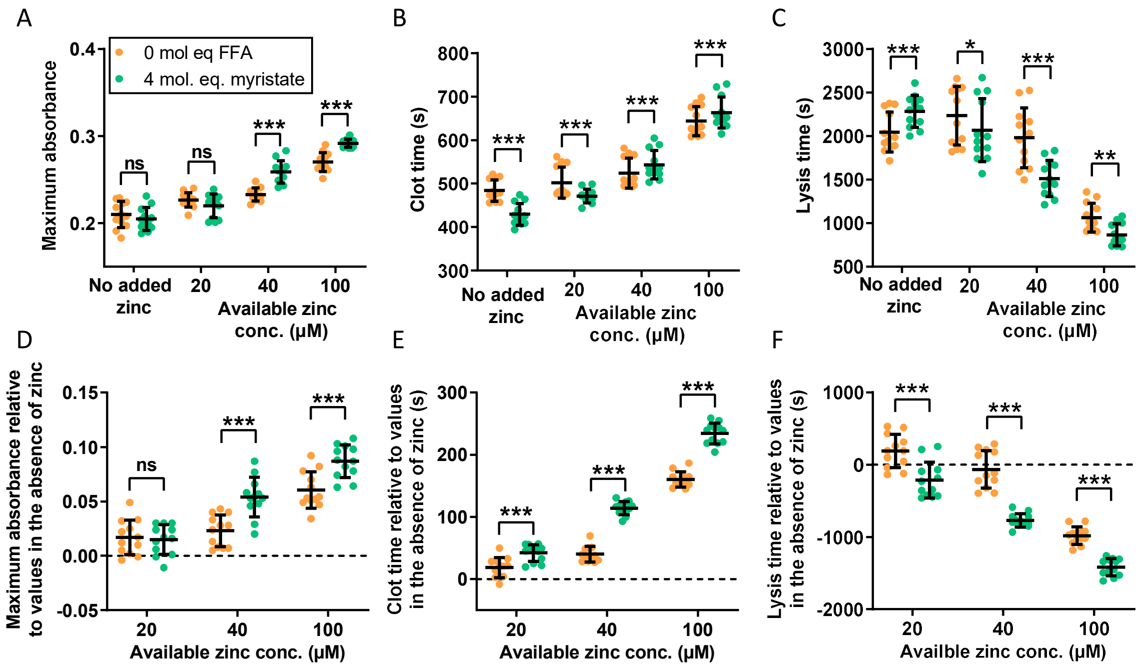


**Figure S28. Effects of $Zn^{2+}$ and FFAs on fibrin clot parameters in pooled plasma and effects relative to the parameter values in the absence of $Zn^{2+}$.** Turbidimetric fibrin clotting and lysis assays were performed in pooled plasma diluted 6-fold in buffer (50 mM Tris, 100 mM NaCl, pH 7.4), with final concentrations of 7.5 mM $CaCl_2$, 0.03 U/ml thrombin, 20.8 ng/ml tPA, while 0-100 μM available $Zn^{2+}$ was added as $ZnCl_2$ (concentration calculated before the dilution of plasma) as well as either 0 or 4 mol. eq. myristate (n=12). Fibrin clot parameters including (A) maximum absorbance, (B) clot time and (C) lysis time were measured. Two-way ANOVA followed by Sidak's multiple comparisons test were used to analyse the data. All three parameters showed a significant difference with the addition of $Zn^{2+}$ ($p<0.0001$, $p<0.0001$ and $p<0.0001$ respectively) as well as with 4 mol. eq. myristate ($p=0.0034$, $p=0.0002$ and $p<0.0001$ respectively). The parameter values relative to their values in the absence of $Zn^{2+}$ were then calculated: (D) maximum absorbance, (E) clot time and (F) lysis time. All three parameters showed a significant difference with the addition of $Zn^{2+}$ ($p<0.0001$, $p<0.0001$ and $p<0.0001$ respectively) as well as with 4 mol. eq. myristate ($p=0.0013$, $p<0.0001$ and $p<0.0001$ respectively). The data is represented as mean ± SD. Statistical significance is indicated with ns where $p>0.05$, * where $p<0.05$, ** where $p<0.01$ and *** where $p<0.001$.

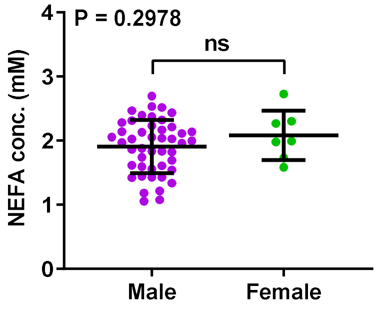


**Figure S29. Comparison of FFA concentrations between sexes in plasma samples from patient with T2DM and controls.** No difference was found.

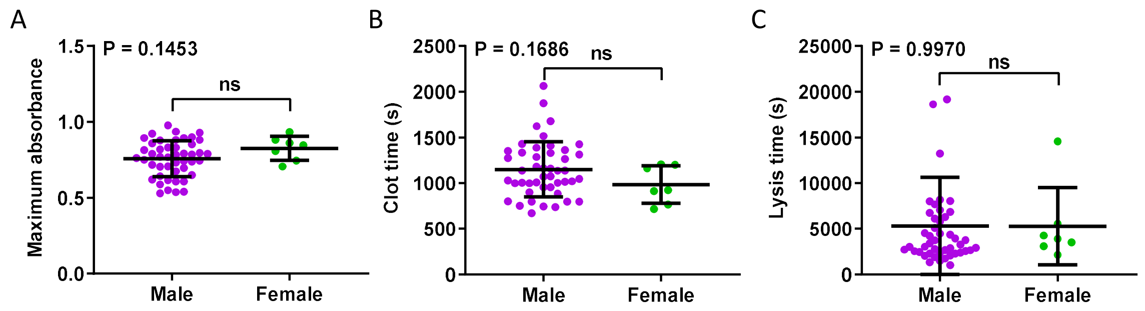


**Figure S30. Comparison of clotting parameters between sexes in plasma samples from patient with T2DM and controls.** (A) Maximum absorbance, (B) clot time and (C) lysis time. No difference was found.

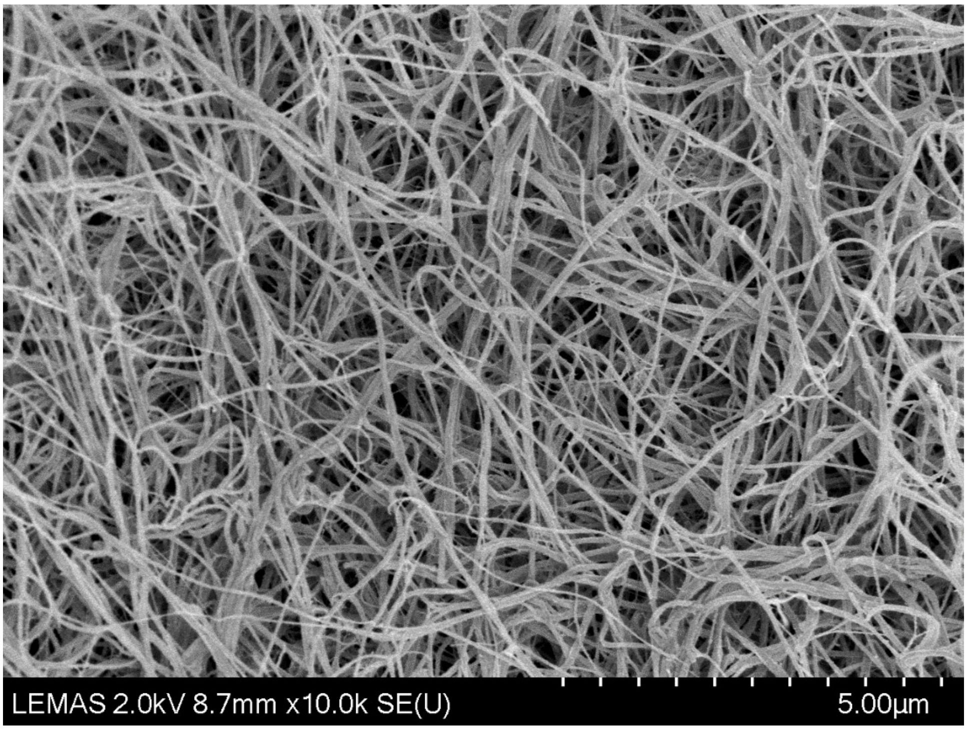


**Figure S31. Representative image from SEM experiments. Purified system, no zinc.** The image was viewed and photographed at ×10,000 magnification using a SU8230 SEM.

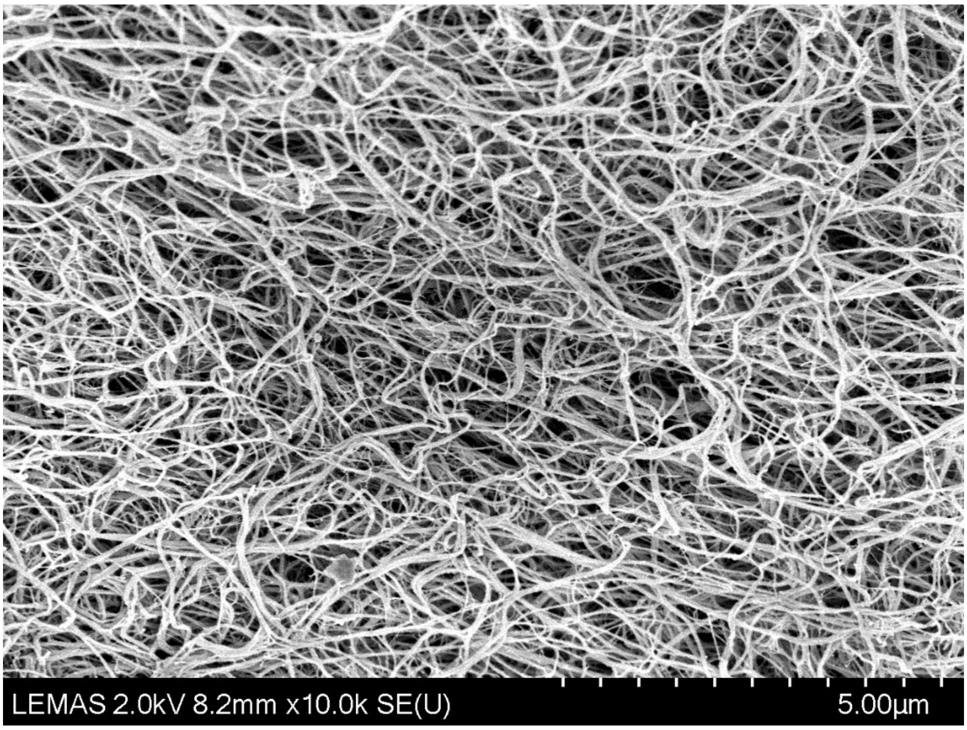


**Figure S32. Representative image from SEM experiments. Purified system, 20 µM zinc.** The image was viewed and photographed at ×10,000 magnification using a SU8230 SEM.

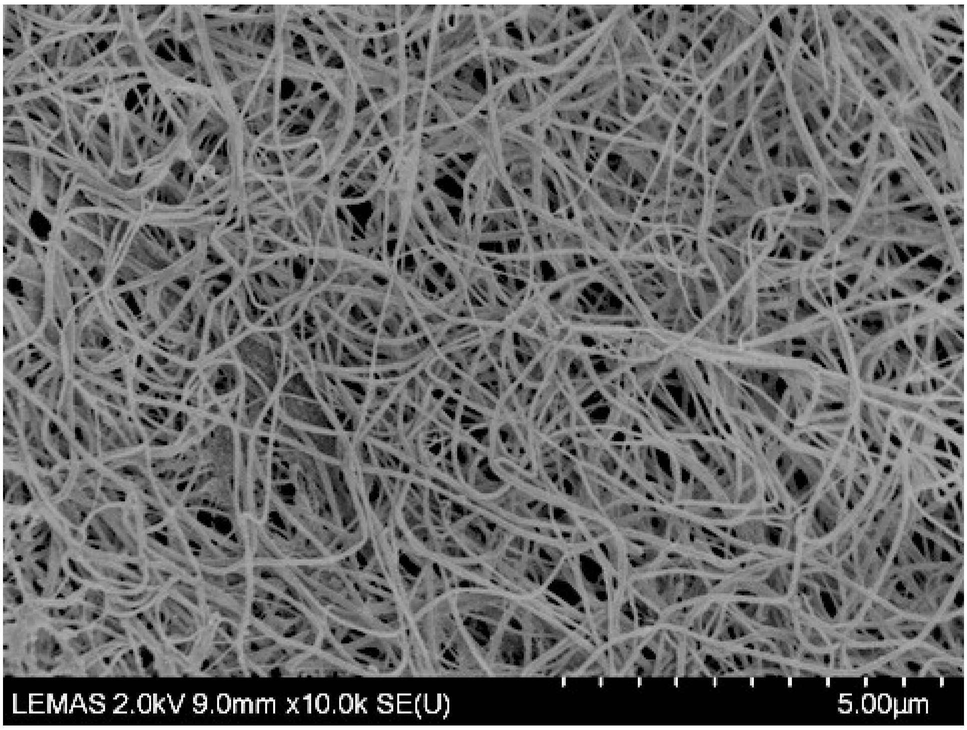


**Figure S33. Representative image from SEM experiments. Controls, no zinc.** The image was viewed and photographed at ×10,000 magnification using a SU8230 SEM.

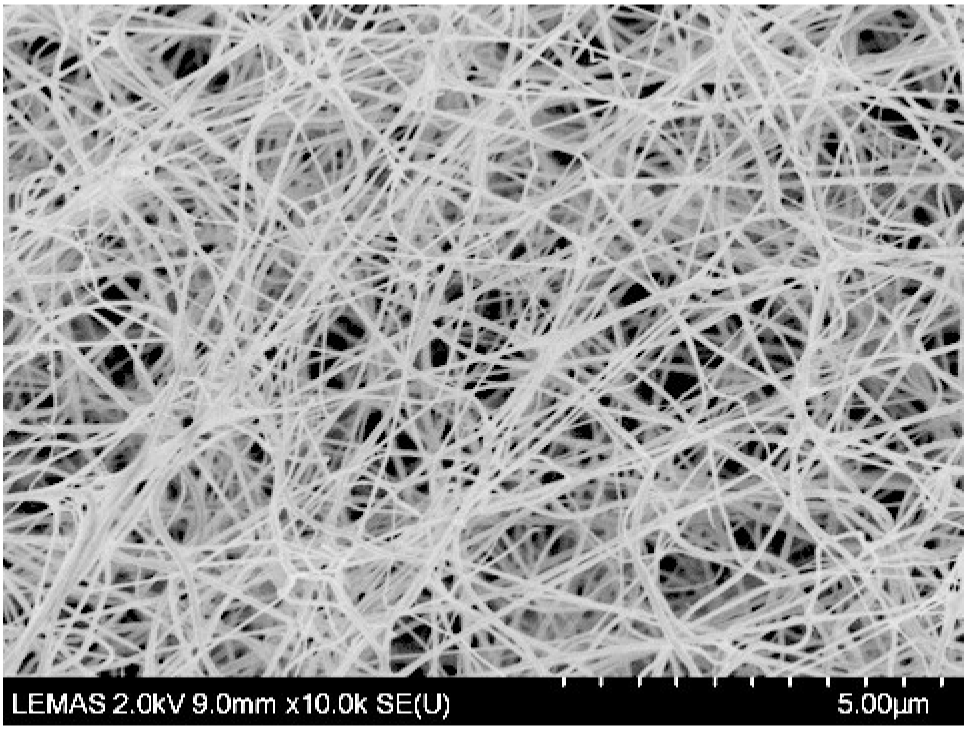


**Figure S34. Representative image from SEM experiments. Controls, 20 μM zinc.** The image was viewed and photographed at ×10,000 magnification using a SU8230 SEM.

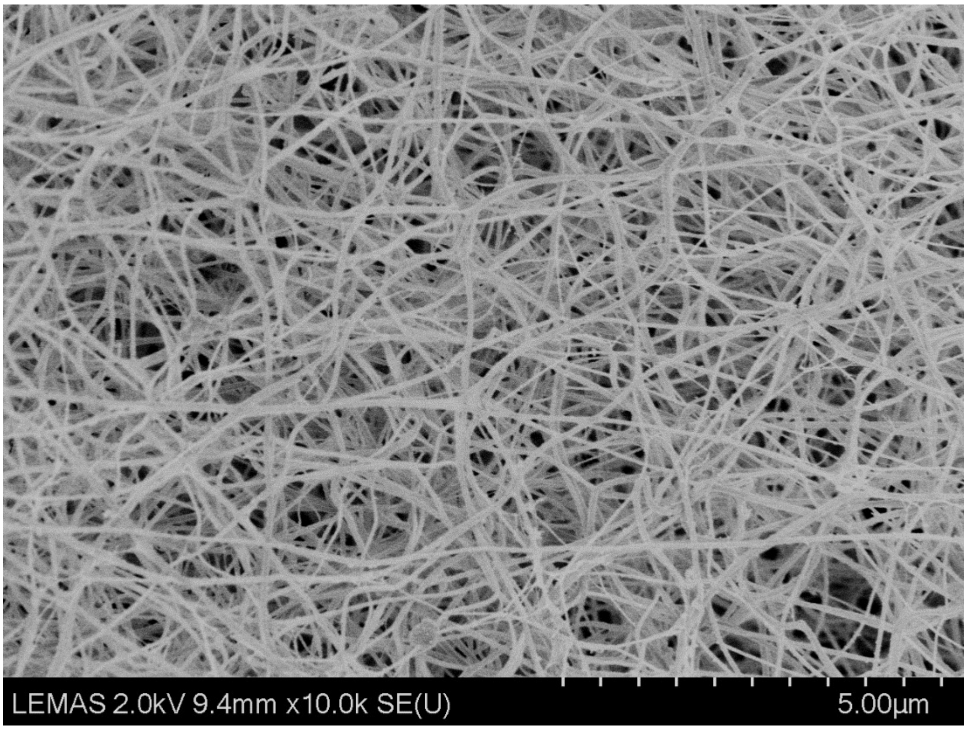


**Figure S35. Representative image from SEM experiments. T2DM, no zinc**. The image was viewed and photographed at ×10,000 magnification using a SU8230 SEM.

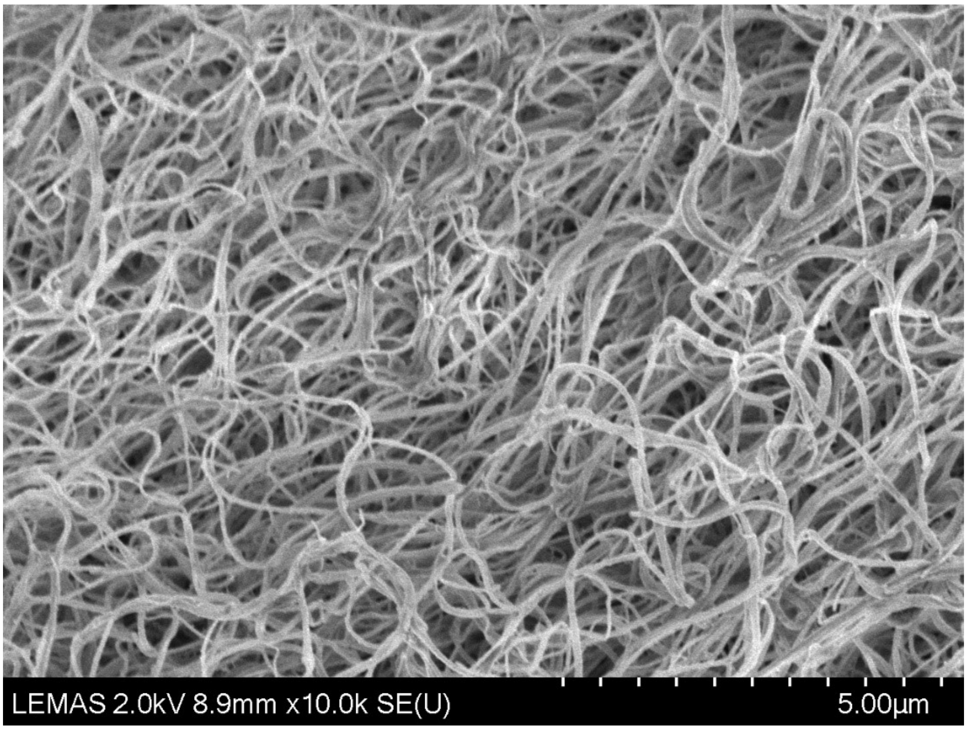


**Figure S36. Representative image from SEM experiments. T2DM, 20 µM zinc.** The image was viewed and photographed at ×10,000 magnification using a SU8230 SEM.
